# Epigenetic Subtypes of High-Grade T1 Bladder Cancer Reveal Intra-Tumor Heterogeneity and Distinct Interactions with Tumor Microenvironment

**DOI:** 10.1101/2025.06.20.659124

**Authors:** Joaquim Bellmunt, Yingtian Xie, Nuria Juanpere, Miguel Gomez Munoz, Shweta Kukreja, Sonsoles Liria Veiga, Rong Li, Xintao Qiu, Yijia Jiang, Alba Font-Tello, Marie Nunez Duarte, Ilana Epstein, Silvia Hernández-Llodrà, Marta Lorenzo, Silvia Menendez, Toni K. Choueiri, Myles Brown, Henry W Long, Paloma Cejas

## Abstract

High-Grade T1 (HGT1) Non-Muscle Invasive Bladder Cancer (NMIBC) is a clinically heterogeneous disease, characterized by unpredictable treatment responses and limited tools for recurrence prediction. Although molecular classification efforts have been made, patient stratification still primarily relies on clinicopathological features, which offer limited clinical precision. In this study, we integrated chromatin profiling in bulk with single-nuclei(sn) RNA-seq, immunohistochemistry and spatial transcriptomics to define epigenetic subtypes of HGT1, characterize their heterogeneity, and investigate tumor–microenvironment interactions. Our findings reveal distinct chromatin profiles differentiating urothelial (URO) and micropapillary (MP) histological variants of high-risk HGT1. We identified three epigenetic states: two within the URO group, luminal-like inflammatory (LLI) and basal-like (BL), and a separate signature unique to MP tumors. Single-cell and spatial resolution approaches validated intratumoral heterogeneity and provided insight into subtype-specific microenvironmental contexts. Notably, ∼40% of URO tumors exhibited spatially distinct coexisting LLI and BL components. BL regions showed enrichment for angiogenesis and hypoxia pathways and were preferentially located near vascular stroma, while LLI regions showed to be located at the core of the tumor. MP tumors featured a markedly different microenvironment, characterized by diverse populations of cancer-associated fibroblasts (CAFs) and M2-polarized macrophages intermingled with tumor cells, suggesting a more immunosuppressive niche which could account for their worse clinical outcome. URO tumors, by comparison, showed a more immune-excluded phenotype. These findings provide a detailed molecular and spatial map of bladder cancer heterogeneity, highlighting how distinct epigenetic subtypes and histological variants are associated with tumor architecture and microenvironmental interactions. Our data underscore the need for subtype-specific therapeutic strategies to more effectively address the complexity already existing at HGT1 bladder cancer.

## INTRODUCTION

Non-muscle invasive bladder cancer (NMIBC) is an early stage of bladder cancer that shows heterogeneity in terms of disease outcome. Despite several attempts there is still a lack of well-established and reliable prognostic factors. High-grade T1 (HGT1) NMIBC is the subgroup that has the highest risk of disease recurrence (∼40%) and progression (∼21%) (Martin-Doyle et al. 2015). Currently, the risk of progression in HGT1 cannot be estimated based on classic clinico-pathologic prognostic markers. Several histological subtypes are known to add poor prognostic outcome on HGT1 management. A well known example is the presence of the micropapillary variant that has been associated with adverse pathological features and poor outcome (Amin et al. 1994). The Micropapillary subtype (diagnosed when there is a micropapillary component > 10%) represents approximately 0.6–2.2% of urothelial Tumors (Samaratunga and Delahunt 2012; Sui et al. 2016; Vourganti et al. 2013).

Efforts have been made to establish a molecular classification of bladder cancer, particularly in the muscle-invasive stage (Sjödahl et al. 2017, 2012). Data from these studies have been jointly reviewed to develop a consensus classification in six subtypes of MIBC with different degrees of luminal and basal characteristics (Kamoun et al. 2020). More recently, molecular classification has been described for NMIBC (de Jong et al. 2023; Hedegaard et al. 2016; Kamoun et al. 2020). The larger NMIBC analysis of the multi-institutional UROMOL study subclassified the tumors into four types (Sia Viborg Lindskrog et al. 2021)(Prip et al. 2025). However, these subtypes are still not being used in clinics. One initial limitation is the extensive intratumor heterogeneity recently described with the advent of the single cell methodologies (Sia V. Lindskrog et al. 2023a; Jin et al. 2023).

An important characteristic of bladder cancer across stages is the occurrence of recurring mutations, some of them affecting genes involved in chromatin structure such as KMT2A and KDM6A (Bellmunt et al. 2020; Robertson et al. 2017). We have previously investigated cancer cell-intrinsic genetics of HGT1 NMIBC by leveraging human patient samples (Bellmunt et al. 2020; Bowden et al. 2020). Our results showed a higher similarity in the pattern of somatic mutations between HGT1 and MIBC as compared to the low-grade (LG) NMIBC (Bellmunt et al. 2020; Kim et al. 2019; Robertson et al. 2018). The genetic similarity between HGT1 and MIBC suggests activation of mechanisms of invasion already in HGT1. Taking the high abundance of genetic alterations in chromatin genes at a similar level between MIBC and HGT1, leads to the hypothesis that the mechanisms associated with progression could be impacted by the chromatin structure (Cejas et al. 2019; Bellmunt et al. 2020). Some of the identified mutations in chromatin genes in HGT1 affect genes involved in different levels of enhancer regulation. For instance, ARID1A, associated with disease progression in HGT1 ((Bellmunt et al. 2020)) is a member of the chromatin remodeling complex SWI/SNF (Mullen et al. 2021), KMT2D methylates H3K4 at promoters and enhancers (Hoffmann and Schulz 2021) and EP300 is a histone acetyltransferase involved in enhancer activation (Robertson et al. 2017; Goodman and Smolik 2000). This high representation of mutations in chromatin remodelers suggests an impact of chromatin reprogramming in BC tumorigenesis across stages.

Chromatin analysis, even in bulk, can reveal truncal lineages because chromatin states are less transient and reflect regulatory potential rather than just current gene expression levels (Cejas et al. 2019; Chung et al. 2019). Based on this, we started performing H3K27ac profiling of HGT1 tumors. This led to the discovery of three chromatin subclasses, two sub-classifying the classic URO histology, and a distinct one associated with the micropapillary (MP) histological subtype. The subtypes differ in relevant molecular characteristics and show association with outcome. Single-cell, immunohistochemistry and spatial transcriptional analyses show that the two URO subtypes are not homogeneous across cancer areas but occupy distinct spatial regions. The differences between URO and MP histologies rely on a markedly distinct tumor microenvironment (TME). MP is characterized by an increased stromal infiltration, enriched populations of cancer-associated fibroblasts (CAFs), and a predominance of M2-polarized macrophages interspersed within the tumor. Our findings highlight the complex heterogeneity of HGT1 bladder cancer and underscore the relevance of single-cell and spatial resolution analyses to advance target discovery and biomarker identification for precision medicine.

## RESULTS

### Chromatin analysis reveals subtypes in HGT1 bladder cancer

We explored the status of activation of enhancers and super-enhancers (SE) by H3K27ac profiling of a cohort of 17 HGT1 patients with clinically annotated outcomes using FiTAc-seq analysis on FFPE archived clinical tissues, as previously described by our group (Font-Tello et al. 2020). We included a set of samples enriched for micropapillary (MP) content, which is known to be associated with a worse clinical outcome (Willis et al. 2015). For all the cases included in the analysis, we ensured a minimum of 80% enrichment in cancer cells, performing macrodissection whenever required. The analysis produced high-quality results in terms of identified peaks and fraction of reads in peaks (summarized in Table 1). The results showed activation of enhancers at previously known genes related to bladder cancer such as *NECTIN4* and *SOX4* across the cases (Fig 1a), validating the specificity and quality of the results.

**Figure 1.**
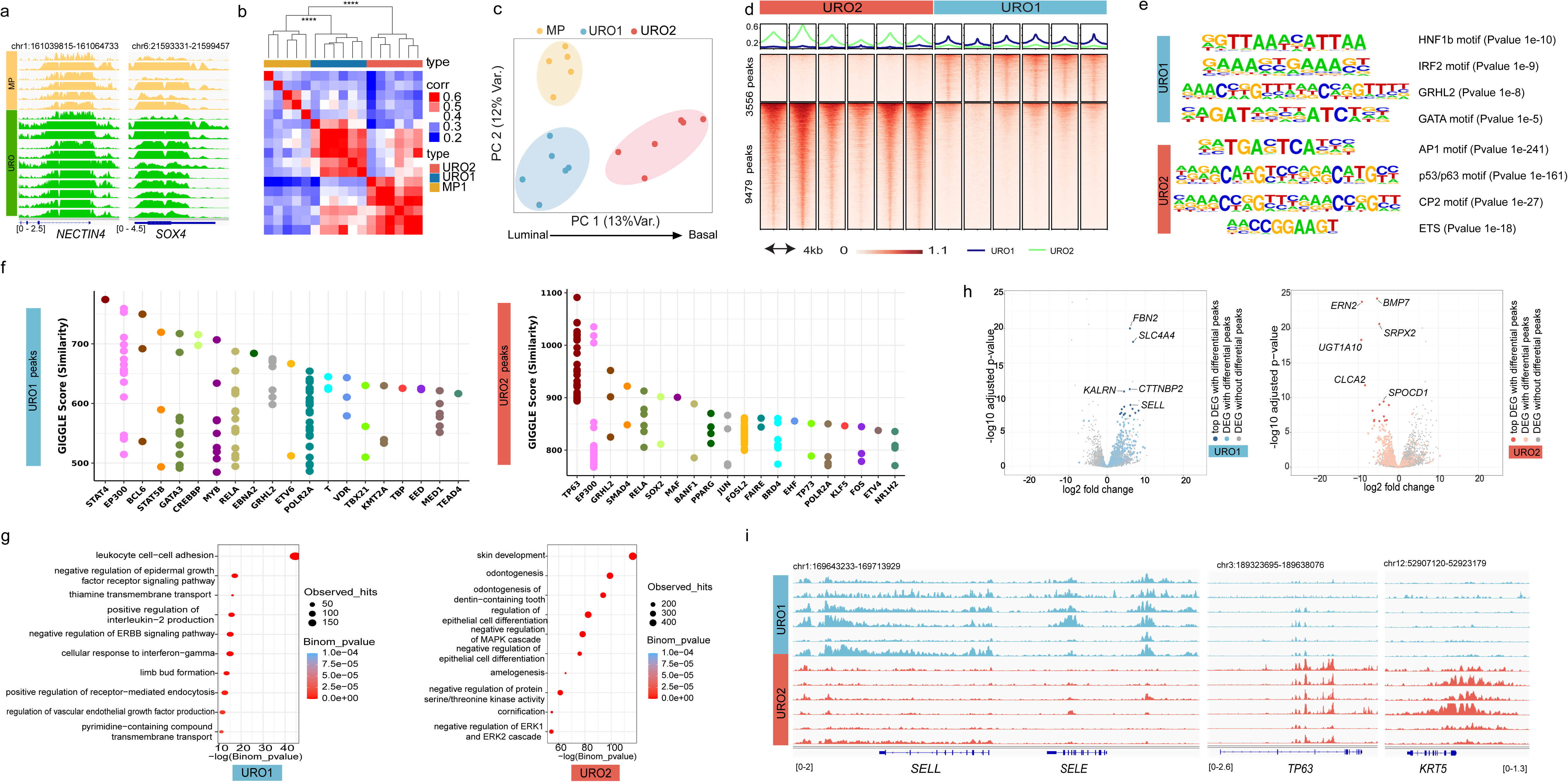
Epigenetic analysis of HGT1 tumors. **a**) Genome tracks of H3K27ac FiTAc-seq data from HGT1 samples at the NECTIN4 and SOX4 loci. **b)** Heatmap showing the pairwise Pearson’s correlation between HGT1 samples. Above the heartmap is hierarchical clustering of the sample correlations indicating the statistical significance of the three main branches of the tree (Pvalue=0.001, Permanova test). **c)** Principal component analysis (PCA) of the H3K27ac data showing the three clusters identified in the correlation analysis. PC1 represents the axis of Luminal to Basal characteristics. **d)** Heatmap representation of the differential H3K27ac regions distinguishing URO1 and URO2 samples. **e)** Results from motif analysis of the differential peaks between URO1 and URO2, showing the top most enriched motif for URO1 (top) and URO2 (botton). **f)** Similarity (GIGGLE) scores between regions of increased H3K27ac signal genome-wide in URO1 (left) and URO2 subtype (right) and public ChIP-seq datasets transcription factors. On the URO1 side, the top overlapping TF is STAT4 in lymphocytes. For the URO2 peaks, the top overlapping TF is TP63 in keratinocytes. **g)** Gene Ontology enrichment results showing the pathways enriched in genes with nearby LLI specific H3K27ac regions (left) and BL specific H3K27ac regions, determined by GREAT analysis. **h)** Association between differential H3K27ac regions and differential gene expression for URO1 (left) and URO2 (right). Each volcano plot depicts RNA-seq log2-fold change (x-axis) and p-value adjusted for multiple hypothesis testing (y-axis) as calculated by DESeq2. Each dot represents one gene, with blue dots (left) indicating significant differentially expressed genes (DEG) associated with a differential H3K27ac region nearby for the LLI subtype. Orange dots on the right, represent DEGs associated with a differential H3K27ac region nearby for the BL subtype. Gray dot: DEGs with no significant differential H3K27ac region nearby. **i)** (Left) Representative IGV tracks at SELL and SELE showing super enhancers for the LLI subtype. (Right) Representative IGV tracks at TP63 and KRT5 super enhancers for the BL subtype.

**TABLE 1.**
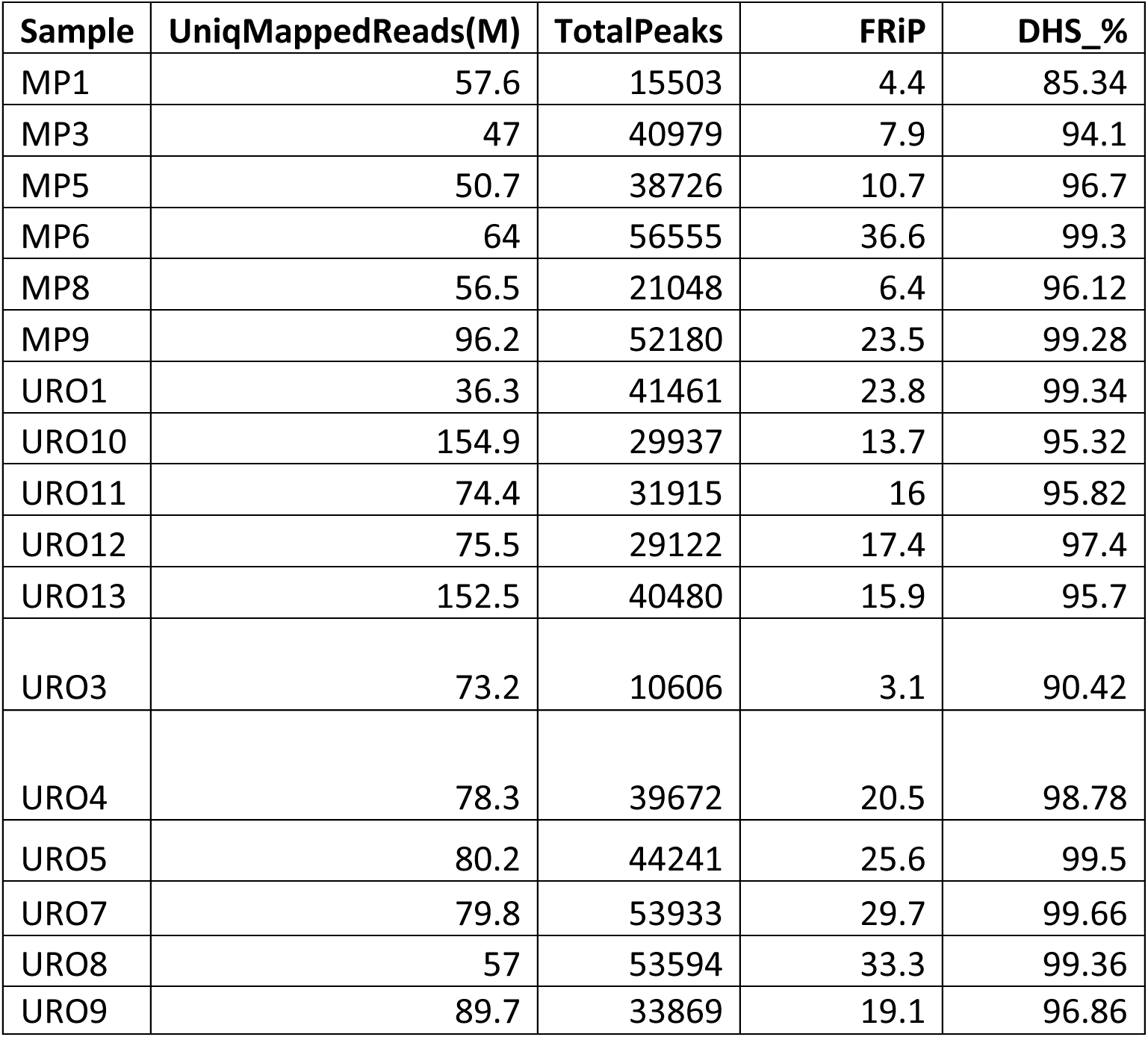
ChIP-seq QC statistics.

| **Sample** | **UniqMappedReads(M)** | **TotalPeaks** | **FRiP** | **DHS_%** |
| --- | --- | --- | --- | --- |
| MP1 | 57.6 | 15503 | 4.4 | 85.34 |
| MP3 | 47 | 40979 | 7.9 | 94.1 |
| MP5 | 50.7 | 38726 | 10.7 | 96.7 |
| MP6 | 64 | 56555 | 36.6 | 99.3 |
| MP8 | 56.5 | 21048 | 6.4 | 96.12 |
| MP9 | 96.2 | 52180 | 23.5 | 99.28 |
| URO1 | 36.3 | 41461 | 23.8 | 99.34 |
| URO10 | 154.9 | 29937 | 13.7 | 95.32 |
| URO11 | 74.4 | 31915 | 16 | 95.82 |
| URO12 | 75.5 | 29122 | 17.4 | 97.4 |
| URO13 | 152.5 | 40480 | 15.9 | 95.7 |
| URO3 | 73.2 | 10606 | 3.1 | 90.42 |
| URO4 | 78.3 | 39672 | 20.5 | 98.78 |
| URO5 | 80.2 | 44241 | 25.6 | 99.5 |
| URO7 | 79.8 | 53933 | 29.7 | 99.66 |
| URO8 | 57 | 53594 | 33.3 | 99.36 |
| URO9 | 89.7 | 33869 | 19.1 | 96.86 |

The unsupervised hierarchical analysis of the H3K27ac-marked (transcriptionally active) regulatory regions (promoter and enhancers) distinguished three statistically different clusters (Fig 1b). Two clusters subdivided the conventional urothelial histology (URO) and the other, corresponded to the MP histologies (Fig 1b,c). Of the two URO clusters, URO1 was closer to the micropapillary (MP) than to the other URO cluster (URO2) (Fig 1c). We next performed differential analysis (Qiu et al. 2021) to identify regulatory sites specific to each of the three clusters (Suppl. Fig 1a). To investigate the transcriptional mechanisms underlying the chromatin differences, we applied HOMER analysis (Huppert et al. 2009) that assessed the enrichment of transcription factors (TF) DNA-binding motifs at the differential regions. The results validated the luminal origin of the MP cluster by showing enrichment in the GRHL2 motif on the differential regions for the MP cluster (Suppl. Fig 1b) (Hoffmann and Schulz 2021; Jovanović et al. 2023). On the other side, the differential regions for the URO2 cluster showed TP63 as the top enriched motif (Table 2), suggesting more basal-like characteristics for this cluster (Hoadley et al. 2014; Murray-Zmijewski, Lane, and Bourdon 2006). Closer to the MP than to URO2, the URO1 cluster showed enrichment in motifs for inflammatory related TFs such as IRF2 and HNF1, supporting a more luminal-like-inflammatory phenotype (Table 2). In summary, the chromatin analysis shows three distinct clusters that are positioned in an axis (PC1) from the luminal to the basal phenotype (Fig 1c).

**TABLE 2.**
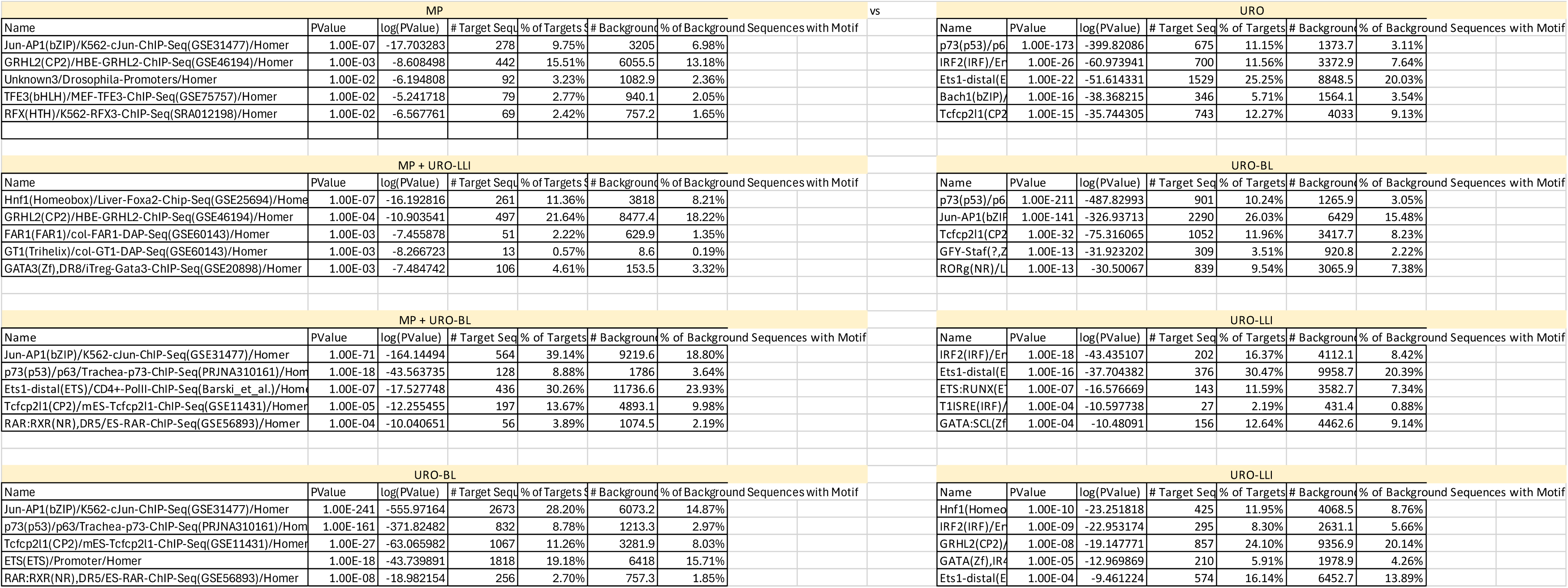
Motif analysis results.

We further investigated the urothelial subtypes without the potential confounding parameter of the MP histology, focusing on the URO cases and assessing the differences between their two clusters. The comparison of URO1 with URO2 resulted in the identification of 3556 regulatory regions differentially H3K27ac-activated in URO1 and 9479 differentially activated in URO2 (Fig 1d) showing differences within the same histological subtype. Motif analysis of these two sets of regions showed similar results to what was observed in the previous analysis, although it included enrichment in GATA motifs on the luminal-like side (Fig 1e). We next compared the differentially active regions to published chromatin immunoprecipitation sequencing (ChIP-seq) profiles compiled in CistromeDB (Taing et al. 2024). The results showed that top URO1-peaks overlap with STAT4 in lymphocyte cells. Also, we found a high overlap with GATA3 binding sites in lymphocytes and epithelial cells. The overlap with these transcription factors binding sites, validates the inflammatory and luminal phenotype revealed by the motif analysis (Fig 1f). The URO2-activated regions showed the highest overlap with TP63 bindings analyzed in keratinocytes validating the basal characteristics (Fig 1f). Consistent subtypes characteristics were also exhibited when differential regions were analyzed by Genomic Regions Enrichment of Annotations Tool (GREAT) analysis (McLean et al. 2010). This tool associates genomic regions with nearby genes and then examines the enrichment of Gene Ontology (GO) pathways for the set of genes associated with each specific subtype. The results showed the immune pathway “Leukocyte cell-cell adhesion” as the top enriched pathway for the URO1 cluster (Fig 1g) while showing enrichment for “skin development” and “cornification” for the URO2 cluster (Fig 1g).

We next integrated the differentially activated promoters and enhancers with gene expression from a subset of the 17 cases for which we had available RNA-seq (Bowden et al. 2020) (Fig 1h) by combining the differential regions between URO1 and URO2 with the corresponding differential gene expression. As shown by the volcano plot, the top differentially expressed genes with corresponding differential enhancer activation in their vicinity, included relevant genes involved in EMT and angiogenesis such as *BMP7, ERN2*, and *SRPX2* (Liu, Fan, and Wu 2017) (M. Jiang et al. 2015) for the URO2 cluster. (Fig 1h). The URO1 side showed a more complex phenotype including a number of genes associated with immune pathways like *SLC4A4* (Cappellesso et al. 2022), *KALRN* (M. Li et al. 2020) and *SELL* (Watson et al. 2019) (Fig 1h). The relevance of the inflammatory genes for the URO1 was further revealed by the presence of Superenhancers (SE) at some of these genes as is the case for *SELL* and *SELE* (Fig 1i) along with SE at luminal genes like *GATA3* (Kouros-Mehr et al. 2006; Warrick et al. 2016) (Suppl. Fig 1c). Conversely, URO2 showed activation of a transcriptional circuit for *TP63* where the motif enrichment is accompanied by the activation of the super-enhancer at the *TP63* locus (Fig 1i, Suppl. Fig 1c) potentially involved in the maintenance of the basal lineage for the URO2 cluster (Saint-André et al. 2016). The basal lineage is further reflected by the existence of a SE at KRT5 (Fig 1i).

Overall, our findings show the presence of distinct chromatin subtypes in HGT1 bladder cancer samples. We showed urothelial histology can be further divided into a basal-like (BL) and a luminal-like inflammatory (LLI) subtype and that MP histology is associated with a distinct chromatin profile. While luminal and basal subtypes have been previously described in bladder cancer, our study establishes a chromatin-based framework for those subtypes and extends it to the MP histological variant.

We next aimed to assess the homogeneity of chromatin subtypes in HGT1 by performing single-cell (sc) ATAC-seq analysis on two independent HGT1 patient samples from a separate cohort (HGT1_1 and HGT1_2). Using our previously established protocol (Cejas and Long 2022), we isolated nuclei from frozen clinical tissue and performed scATAC-seq sequencing with 10X Genomics. After filtering out low-quality nuclei, we retained 12,173 nuclei across both samples for analysis (Suppl. Fig. 1d). A combined UMAP analysis of the two patient samples revealed multiple clusters (Suppl. Fig. 1d). To distinguish tumor from normal cell populations, we applied Copy Number Variation (CNV) inference (Nikolic et al. 2021; Ramakrishnan et al. 2023) (Suppl. Fig 1e). Notably, while clusters of normal cells overlapped between the two patients, the clusters of cancer cells remained distinct, indicating greater chromatin structural differences in tumor cells compared to normal cells (Suppl. Fig. 1d, 1f). To further characterize normal cell lineages, we leveraged an atlas of previously published scATAC-seq datasets from normal human tissues (Y. Jiang et al. 2024) By projecting our samples onto the UMAP representation of this reference dataset, we identified a diverse set of normal cell types, including B cells, T cells, macrophages, endothelial cells, and smooth muscle cells (Suppl. Fig. 1f).

The top genes with differentially activated enhancers and corresponding overexpression in the bulk analysis were used to generate two 15-gene signatures, designed to classify cancer cells as either LLI or BL subtypes (Fig. 1h) (from now on: Chromatin derived score (CDS) (Table 3). To further investigate subtype distribution, we applied the gene activity score from this signature to the scATAC-seq data (Suppl. Fig. 1g). The analysis revealed distinct differences between the two tumors: HGT1_1 exhibited a stronger enrichment for the BL subtype, whereas HGT1_2 was more enriched for the LLI subtype (Suppl. Fig. 1g).

**TABLE 3.** Chromatin derived gene signatures.

| **LLI** | **BL** |
| --- | --- |
| FBN2 | BMP7 |
| SLC4A4 | ERN2 |
| CTTNBP2 | SRPX2 |
| KALRN | UGT1A10 |
| SELL | CLCA2 |
| PMP22 | SPOCD1 |
| UGT2B7 | SERPINB5 |
| COL12A1 | SEMA4B |
| PTH2R | CLCA4 |
| ANKRD36 | CLU |
| DCDC2 | UGT1A7 |
| PRTG | SH3PXD2A |
| BPGM | AQP3 |
| PDE9A | ANXA10 |
| ANXA3 | SLITRK6 |

Supporting these classifications, the subtypes aligned with well-established lineage markers. BL-scored cancer cells showed increased KRT5 promoter accessibility along with a higher enrichment of the TP63 motif (Suppl. Fig. 1h, i). Conversely, cells with lower BL scores exhibited increased KRT20 promoter accessibility and enrichment of GATA3 motifs, consistent with an LLI phenotype (Suppl. Fig. 1h, i). Despite an overall homogeneity within each tumor, the alignment between CDS scores and known markers was not complete. A subset of cells lacked accessibility at both keratin promoters, indicative of terminal differentiation states (Suppl. Fig. 1g). This incomplete overlap suggests a spectrum of differentiation within each subtype, with only a fraction of cells expressing terminal markers. Such a continuum of differentiation states could also explain the varying enrichment of transcription factor motifs observed intratumorally across clusters (Suppl. Fig. 1j).

Overall, we concluded that the CDS classifier, derived from a chromatin-based signature, may provide a robust lineage stratification independent of the differentiation status.

### Expression of the LLI and BL chromatin derived subtypes in two independent urothelial bladder cancer patient cohorts

To assess the representation of the two URO subtypes, Basal-Like (BL) and Luminal-Like Inflammatory (LLI), in an expanded cohort of high-grade T1 (HGT1) urothelial tumors, we applied the Chromatin Differentiation Score (CDS) to score a bulk RNA-seq dataset of 62 previously collected and analyzed HGT1 cases (Bowden et al. 2020) The CDS results revealed a continuous distribution of scores, ranging from cases with high BL signature to those with high LLI signature, with many cases exhibiting mixed characteristics (Fig. 2a). A similar spectrum was observed in the UROMOL dataset, an independent cohort of 438 NMIBC cases including 78 HGT1 tumors (Sia Viborg Lindskrog et al. 2021) (Fig. 2b). In both cohorts, LLI cases exhibited significantly higher expression of *KRT20*, while BL cases were enriched for *KRT5* and *TP63* expression (Fig. 2c). Interestingly, other luminal markers such as *PPARG*, *GATA3*, and *CDH3* did not show subtype-specific expression patterns in any of the two cohorts (Fig. 2c, Suppl. Fig. 2a, b), potentially due to lower expression levels and higher inter-patient variability compared to keratin genes.

**Figure 2.**
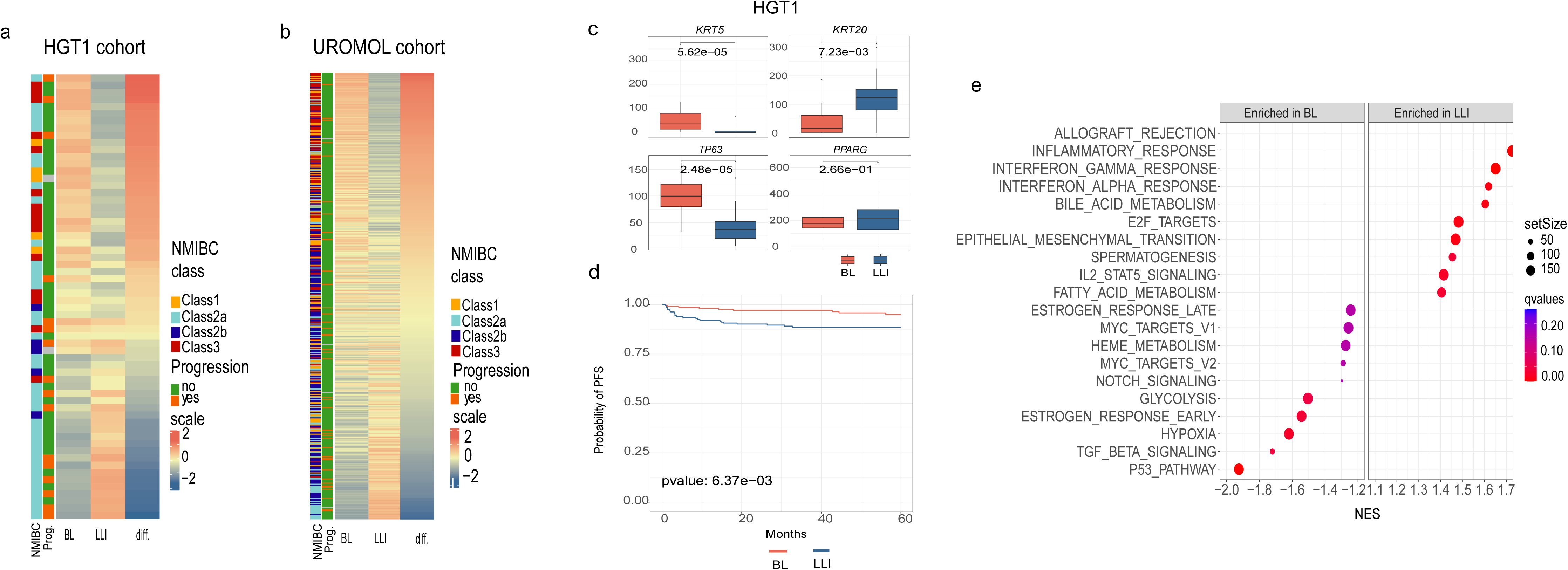
Analysis of chromatin-derived signatures in two NMIBC patient cohorts. **a**) Heatmap representation of BL and LLI signature scores of the HGT1 gene expression cohort. Patients are ordered from most BL to most LLI and also annotated by UROMOL2021 classification into 4 subclasses (1, 2a, 2b and 3). The clinical outcome is also annotated for the cases as cancer progression (yes and no). **b**) Heatmap representation of BL and LLI signature scores of the UROMOL gene expression cohort, ordered and annotated as in **a**. **c)** Boxplots comparing the expression of marker genes analyzed in cases scored as BL (red) or LLI (blue) in the HGT1 cohort. *KRT5*, *KRT20* and *TP63* show statistically differences between sub groups (difference analyzed by t-test). **d)** Kaplan-Meier survival curves for BL (red) and LLI (blue) groups in the UROMOL cohort. Cases were dichotomized between BL and LLI based on the median of the difference in score values. **e)** Gene Set Enrichment Analysis (GSEA) of BL and LLI differential genes in the HGT1 cohort. The differential analysis was done comparing the top quartiles of score values (top BL vs. top LLI).

To contextualize our classification, we compared our CDS-based subtypes with the molecular subtypes described by Lindskrog et al (UROMOL2021), which define four NMIBC classes with varying basal and luminal features (Sia Viborg Lindskrog et al. 2021). While our classification defines two distinct chromatin-defined subtypes, notable similarities emerged. For instance, BL and UROMOL2021 Classes 1 and 3 shared elevated *TP63* expression, while LLI and UROMOL2021 Class 2a were both characterized by high *KRT20* and *ERBB2* expression (Fig. 2a, b). These overlapping molecular features support the existence of biologically distinct NMIBC subgroups with chromatin-associated differences.

We observed a significant overlap between the LLI subtype and UROMOL2021 Class 2a in the UROMOL cohort (Chi-squared test, p=3.3E-08), while BL showed features in common with Class1 and showed significant overlap with Class 3 (Chi-squared test, p<2.2 E-16). Since UROMOL2021 Class 2a has been linked to increased risk of disease progression (Sia Viborg Lindskrog et al. 2021), We next aimed to stratify UROMOL cases by CDS (BL vs. LLI) based on the median score to evaluate LLI and patient risk. Our result also revealed that the LLI subtype was significantly associated with higher risk of progression (Fig. 2d). This trend was also seen in the 62-HGT1 cohort, where LLI cases had a significantly increased likelihood of progression to muscle-invasive bladder cancer (MIBC) (p = 0.02; Fig. 2a), despite the smaller sample size (Fig. 2a).

To further explore the biological differences between the subtypes, we performed differential gene expression analysis between the top and bottom CDS quartiles in the 62-HGT1 cohort (Fig. 2a), identifying 1,115 differentially expressed genes. Gene Set Enrichment Analysis (GSEA; Subramanian et al., 2005) revealed that LLI tumors were enriched in inflammatory pathways, including “Allograft rejection,” while BL tumors were enriched for “p53 pathway”, “TGF_BETA signalling” and “Hypoxia” (Fig. 2e).

Taken together, these findings confirm the existence of two distinct chromatin states, BL and LLI subtypes, in clinical HGT1 bladder cancer samples from two independent cohorts. The results validate their associated transcriptional programs and shows that the LLI subtype is linked to poorer clinical outcomes.

### Single cell expression analysis validates the existence of subtypes and shows intratumor heterogeneity

To assess heterogeneity at a single-cell resolution in a larger cohort, we performed single-nuclei RNA sequencing (snRNA-seq) using GEM-X Flex technology from 10XGenomics, on nine FFPE tumors from the 62-HGT1 cohort, along with two additional MIBC surgical specimens. These later tumors were selected based on their combined micropapillary >30% and urothelial carcinoma (URO) histology, (hereafter referred to as “MPBC1” and “MPBC2”) which could be validated histologically and compared with the transcriptional findings. Nuclei were isolated using an in-house FFPE-optimized protocol, which builds on our previously published method for frozen tissue (Cejas and Long 2022) and includes additional deparaffinization and rehydration steps.

This approach yielded 37,879 high-quality nuclei across the 11 cases. Data were visualized via UMAP to examine both inter- and intra-tumoral heterogeneity (Suppl. Fig. 3a). Malignant and non-malignant populations were distinguished using a combination of CNV inference via inferCNV (Tirosh et al. 2016) and the expression of cancer markers such as *NECTIN4* (Suppl. Fig. 3b,c). Consistent with our scATAC-seq findings, normal cell clusters were more conserved across patients, whereas tumor cell clusters displayed higher heterogeneity (Suppl. Fig. 3a, 3d). Lineage annotation based on canonical markers identified diverse normal cell types, including T cells, B cells, endothelial cells, fibroblasts, plasma cells and normal epithelial cells (Suppl. Fig. 3d,e). These findings confirm the ability of our FFPE snRNA-seq workflow to preserve TME complexity.

Focusing on the malignant cells, we evaluated cancer subtypes (Fig 3a). To that aim, we extended the CDS framework to include a MP-specific signature, derived analogously to the BL and LLI signatures by identifying MP-specific enhancers and corresponding gene expression (Suppl. Fig. 3f). Then we performed subtype scoring using the CDS scoring, classifying cells as BL, LLI, or MP (Fig. 3b,c). We observed varying degrees of intra-tumoral heterogeneity, with BL and LLI cancer cells coexisting within individual tumors, especially among URO carcinomas (Fig. 3c–d). Reassuringly, the MP signature showed the highest scores in MPBC cases with mixed MP/URO histologies. In these instances, the URO component of MPBC1 scored mainly as BL subtype, whereas in MPBC2, the URO component was classified as LLI (Fig 3d). These findings further suggest that the HGT1-derived signatures could also be applicable to MIBC tumors.

**Figure 3.**
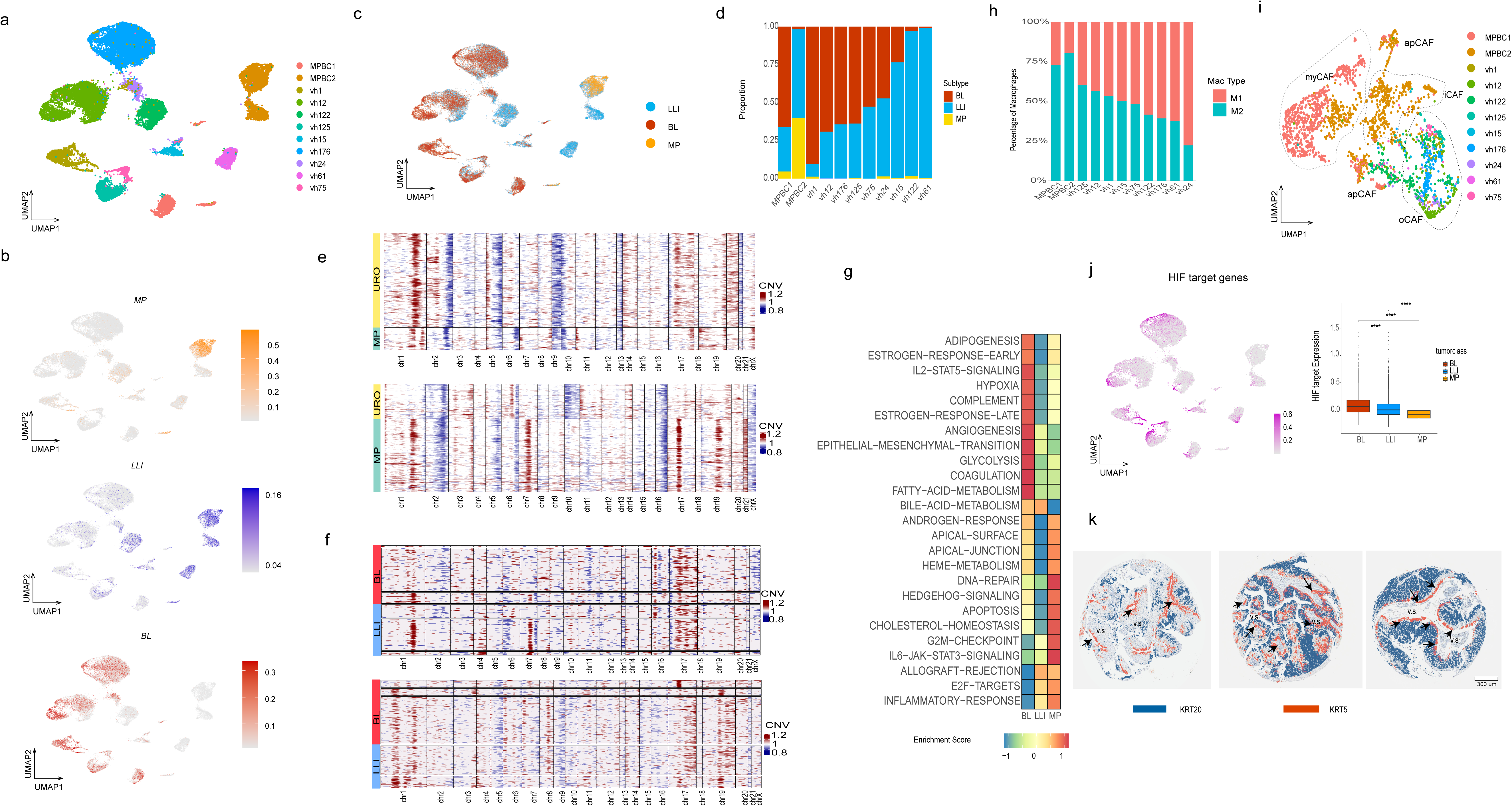
snRNA-seq study of nine HGT1 cases and two MPBC cases with mixed URO and MP histology. **a)** UMAP analysis of the cancer cells from all eleven datasets colored by sample ID. **b)** UMAP analysis showing the scoring of all the cells in our study for the CDS classification system using MP, LLI and BL chromatin derived signatures. **c)** UMAP analysis showing the CDS classification of the cancer cells as LLI (blue), BL (red) or MP subtype (yellow). **d)** Proportional barplot showing the relative percentage of each subtype in each tumor sample. **e)** CNV analysis of the tumor cells in the two MPBC samples clustered by histological subtype. The two histologies (URO and MP) show distinct CNV patterns indicating that they are different clones. **f)** CNV analysis of two representative URO samples (vh75 and vh125) whose cells have been seperated by CDS subtypes and then clustered with K-means to show the subclonal structure. In both cases the BL and the LLI cells show similar clonal structure. **g)** GSEA pathway analysis of the differentially expressed genes for BL, LLI and MP tumor cells across the entire cohort. **h)** Proportional barplot showing the relative percentage of M1 and M2 macrophages in each tumor sample. **i)** UMAP analysis of all CAFs identified across the eleven datasets and colored by sample ID. Dashed lines delimit the populations of canonical CAF subtypes: myCAF, iCAF, and apCAF. CAFs not identified with canonical markers are classified as other (o)CAFs. myCAFs are only represented in the two MPBC samples. **j)** UMAP analysis showing the enrichment of the HIF target gene signature for all the tumor cells across the eleven cases. Box plot (right) shows the quantification of the enrichment across the three CDS subtypes. **k)** Composition of the KRT5 and KRT20 immunostaining for three representative NMIBC cases. The two stainings have been merged showing that the positive cell populations are nearly mutually exclusive. Also, the KRT5-positive cells (orange) are largely near the blood vessels (v.s.) with the KRT20-positive cells (blue) further away.

Interestingly, a few tumors diagnosed as pure URO histology, such as vh1, vh24, and vh122, contained small cell populations with elevated MP scores, suggesting previously unrecognized MP histology (Fig. 3d). Among the URO cases, several tumors: vh12, vh176, vh125, vh75, vh24, and vh15, exhibited a mixture of BL and LLI cells, while vh1 and vh61 were predominantly BL and LLI, respectively (Fig 3d). Tumor vh122, although classified as LLI, lacked KRT20 expression and harbored an MSH2 splice-site mutation (Bellmunt et al. 2020), which may enhance neoantigen production and drive inflammation (Fig. 3c,d). In accordance with their lineage characteristics the snRNA-seq results also showed that BL subtype was associated with *KRT5* and *TP63*, and LLI/MP with *KRT20* (Suppl. Fig. 3g).

We next compared the CNVs profiles across subtypes. In the two mixed-histology cases (MPBC1 and MPBC2), MP and URO components harbored distinct clonal profiles showing two distinct genetic backgrounds (Fig. 3e). In contrast, inside the URO histology, the BL and LLI subtypes shared similar clonal origins (Fig. 3f), suggesting epigenetic regulation rather than genetic divergence. The distinct chromatin subtypes with the same genetic characteristics support the occurrence of cancer cell plasticity inside URO histology, based on chromatin switch. In contrast, the distinct genetic structure of MP and URO histologies suggest more stable subtypes.

To benchmark our classification strategy, we compared CDS scores with the previously proposed UROMOL2021 subtypes (Sia Viborg Lindskrog et al. 2021). When applying the UROMOL2021 classification to the nine HGT1 snRNA-seq datasets, we observed coexistence of multiple subtypes within individual tumors, consistent with previously reported intra-tumor heterogeneity (Sia V. Lindskrog et al. 2023b) (Suppl. Fig. 3h). Notably, we could not find cells classified as class 2b in our single-cell analysis (Suppl. Fig 3h). This finding suggests that this subtype may be the result of a mixed phenotype at the bulk level rather than to a distinct tumor entity which highlights the limitations of bulk transcriptomic profiling for accurate subtype classification.

We then used pseudo-bulk clustering and single-sample GSEA to identify pathway enrichments per subtype (Fig. 3g). MP clusters were enriched in pathways related with metabolism like “cholesterol-homeostasis” and in cell polarity like “apical junction” pathways; LLI clusters in “inflammatory” and “allograft rejection” pathways, and BL clusters in “angiogenesis” and “hypoxia” (Fig. 3g). These pathways involve the interaction between cancer cells and the microenvironment that single cell methodologies enable investigating.

When analyzing the TME, we observed several notable features. One consistent finding across all subtypes was the presence of M2 macrophage polarization, as determined using canonical markers (Fig 3h) (as described in methods). In particular, MP tumors demonstrated activation of mucins, like MUC4 involved in glycosylation, which are known to contribute to a pro-tumorigenic immune environment. For instance, the SPP1–MUC4 axis has been implicated in promoting M2 polarization (Yang et al. 2024) (Elomaa et al. 2025). Consistently, the two MPBC samples with MP components showed a higher abundance of M2-polarized macrophages. However, this observation requires validation in a larger cohort to assess its broader relevance. Overall, the M2 polarization observed in bladder cancer suggests a pervasive, subtype-independent immunosuppressive microenvironment.

We next analyzed the status of Cancer-Associated Fibroblasts (CAFs), which are known to promote M2 macrophage polarization (Monteran and Erez 2019). CAFs can be categorized into three subtypes: Myofibroblastic CAFs (myCAFs), Immune-regulatory Inflammatory CAFs (iCAFs), and Antigen-Presenting CAFs (apCAFs) (Lavie et al. 2022). Previous studies using inferred CAF signatures from the UROMOL bulk RNA-seq cohort associated UROMOL2021 class 2b with high fibroblast signature activity and class 3 with immune depletion and low CAF activity (Caramelo et al. 2023). However, these subtypes do not appear to represent clearly distinct entities when analyzed using single-cell approaches, and the characteristics of CAFs in bladder cancer remain poorly defined.

Our single-cell resolution data provide new insight into this question. All three CAF subtypes were represented in the cohort. Both MPBC samples exhibited myCAF populations, with MPBC2 also showing a mixture of myCAFs and iCAFs (Fig 3i). Notably, VH122, which harbors an MSH2 mutation, was enriched for apCAFs (Fig3i), suggesting a potential role for apCAFs in the neoantigen presentation as a result of MSH2 mutation.

Within the URO histology group, we observed notable differences in pathway enrichment between subtypes (Fig 3g). “Hypoxia” and “angiogenesis” pathways were significantly enriched in the BL subtype compared to LLI, with marked differential activation of the HIF1 target pathway in BL tumors (Fig 3j). These findings suggest that chromatin-defined subtypes are associated with distinct microenvironmental conditions.

To spatially validate these differences, we performed Immunohistochemistry (IHC) on tissue microarrays (TMAs) from an independent cohort of 162 pure urothelial tumors (Lloreta et al. 2017), comprising 102 high-grade T1 (HGT1), 41 low-grade, and 19 muscle-invasive bladder cancer (MIBC) samples. We used KRT5, TP63, and KRT20 as surrogate markers for the BL and LLI subtypes. Notably, coexistence of KRT5- and KRT20-expressing tumor cell populations was seen in 44% of samples (53/120), reflecting intratumoral heterogeneity (Suppl. Fig. 3i). In contrast, exclusive expression of KRT5 (27%), KRT20 (21%), or neither marker (8%) further underscored subtype diversity, patterns consistent with those seen in our snRNA-seq dataset (Fig. 2a, b).

Spatial mapping confirmed subtype-specific cellular organization: KRT5 positive cells were enriched at the tumor–stroma interface, whereas KRT20 positive cells were centrally located (Suppl. Fig. 3j). TP63 overlapped primarily with KRT5, identifying a distinct BL subset. Minimal co-expression supported the statistical spatial separation of these subtypes (Suppl. Fig. 3l, k). The localization of basal KRT5 positive cells significantly nearer the vascular stroma (Suppl. Fig 3k) supports the idea that oxygen availability may play a role in maintaining the BL and LLI phenotypes, with BL cells showing increased sensitivity to hypoxic signaling.

Together, these findings underscore the power of single-cell and spatial transcriptomics, like IHC, to resolve the molecular and spatial complexity of bladder cancer. They highlight the coexistence of transcriptionally distinct subtypes within individual tumors and emphasize the biological relevance of spatial organization for understanding tumor heterogeneity and progression.

### Assessment of subtypes and microenvironment by spatial whole transcriptional analysis

We next applied the 10x Genomics Visium HD platform to spatially investigate the transcriptional profiles of bladder cancer subtypes and visualize their interactions with the TME. This technology enables transcriptome-wide spatial profiling at an enhanced resolution (2 x 2 µm contiguous barcoded squares), and integrates the hematoxylin-stained histological imaging, enabling the spatial mapping of transcription between cancer cells and their surrounding stroma. We conducted this analysis on the two MPBC cases for which we had surgical specimens containing both micropapillary (MP) and urothelial (URO) histologies. Unlike biopsies, these samples preserved the tissue architecture required for spatial transcriptomic profiling (Fig. 4a, b, Sup Fig. 4a). We performed cell segmentation to assign the transcriptional information to each cell by using Bin-to-Cell methodology that combines the H&E image with the transcriptional information, as described in methods. Next, we performed clustering analysis based on a graph-based method (Fig 4a,b and Suppl. Fig 4b).

**Figure 4.**
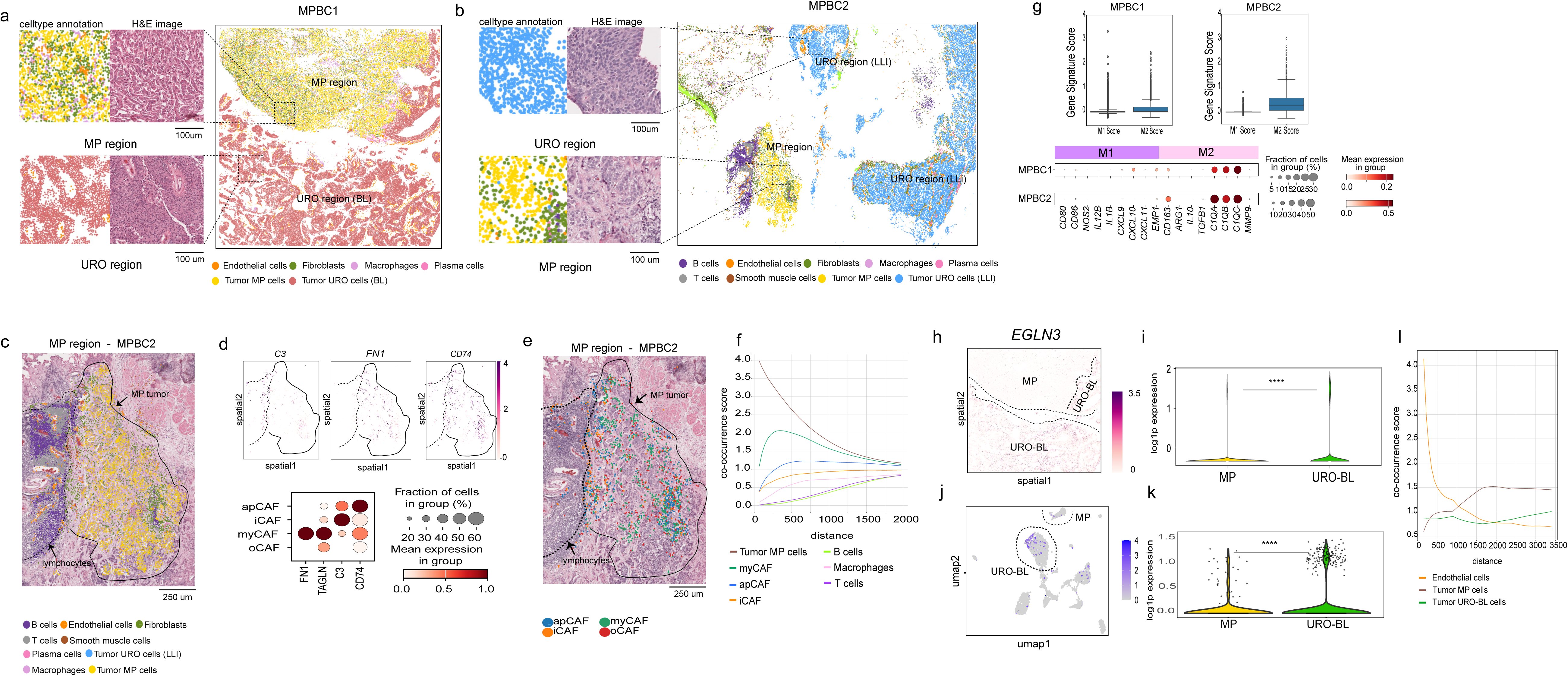
Spatial transcriptomic analysis of two MPBC cases with mixed URO and MP histology. **a)** Spatial transcriptomic results from MPBC1. Dashed boxes highlight regions that are shown in greater detail on the left, where corresponding H&E images and annotated cell types are displayed for representative MP and URO areas. Individual cells are colored by cell type. **b)** Spatial transcriptomic results from MPBC2 displayed in the same way as in **a**. **c)** Enlarged view of the MP region in MPBC2 with cell type annotations overlaid on the H&E image. Solid line surrounds the MP tumor cells and the dashed line demarcates an area enriched in immune cells. **d)** Top panel: spatial representation of canonical CAF markers on the area in **c**. Botton panel: Dot plot visualizing the mean expression of canonical markers to identify the CAF subtypes (apCAF, iCAF, myCAF). **e)** Same region as in **c** showing CAF subtype annotation on top of the H&E image. The solid line surrounds the MP tumor cells and the dashed line demarcates an area enriched in immune cells. **f)** Quantitative analysis of CAF subtype co-occurrence relative to the micropapillary tumor cells in MPBC2. myCAFs show the strongest spatial association with the MP population, exhibiting the highest co-occurrence scores across all distances. apCAFs are more peripherally located, with intermediate co-occurrence. In contrast, iCAFs display minimal co-occurrence and are largely excluded from the tumor region, instead clustering near immune cell-rich areas. **g)** Top panel: Box plots illustrate M1 and M2 gene signature scores derived from canonical markers in MPBC1 (left) and MPBC2 (right). Bottom panel: Dot plot visualizing the expression of canonical M1 and M2 polarization markers for both MPBC samples. **h)** Spatial representation of EGLN3 gene expression in the MP and URO-BL cells of sample MPBC1. **i)** Violin plot of EGLN3 gene expression levels in the MP cells versus the URO-BL cells in the MPBC1 spatial dataset. EGLN3 shows significantly higher expression in the URO-BL compared to MP cells (t-test; p<0.01). **j)** UMAP representation of EGLN3 gene expression in snRNA-seq data from MBPC1. **k)** Violin plot of EGLN3 gene expression levels in the MP cells versus the URO-BL cells in snRNA-seq data from MBPC1. Significantly higher expression is seen in the URO-BL cells (t-test; p<0.01). **l)** Quantitative analysis of endothelial cell co-occurrence with MP and URO tumor regions, based on the area shown in Suppl. Fig 4e. In this tumor, the URO component is entirely BL. These BL cells are more closely associated with endothelial cells than the MP cells, which show no significant spatial enrichment near endothelial cells.

Unsupervised clustering revealed distinct clusters corresponding to normal cell lineages and cancer cell subtypes (Fig. 4a,b, Suppl. Fig 4b). We could assign the normal and tumor clusters to the ones identified by snRNA-seq on those same samples, translating the information from snRNA-seq to a spatial dimension. Notably, MP and URO histologies formed transcriptionally distinct clusters (Suppl. Fig 4b) that spatially matched their corresponding histological spatial locations (Fig 4a, b). In MPBC1, the URO component scored as BL subtype without significant LLI features, differing from snRNA-seq results for the same tumor, likely due to sampling from different tissue sections (Suppl. Fig 4c). Conversely, the URO component of MPBC2 consistently scored as LLI subtype across both platforms (Suppl. Fig 4 d).

Spatial organization of the TME varied significantly between MP and URO components. The URO-associated stroma displayed a relatively simple, immune-excluded phenotype, while the MP histology featured a more complex stromal architecture (Fig 4a, b). In both MPBC1 and MPBC2 (CAFs) and macrophages were found intermixed within the MP region (Fig. 4a, b).

Since our snRNA-seq analysis revealed the presence of distinct CAF populations in the MP samples, we aimed to investigate the spatial location of those cells. We found a particularly complex CAF representation in the MP component of MPBC2 (Fig 4c) that showed presence of myCAF, iCAF and apCAFs based on expression of corresponding canonical markers (Fig 4d). These CAF subtypes occupied distinct spatial niches: myCAFs were embedded within the tumor core, whereas iCAFs localized to the periphery and were interspersed within the cluster of immune cells that included B cells, T cells and macrophages (Fig 4e). We also found populations of apCAF on both sides of the tumor (Fig 4e). We further analyzed the co-occurrence probability of the CAF populations relative to the MP cancer cells. The results validated the spatial observation, showing that myCAF were more likely to co-occur with MP cancer cells (Fig 4f). In MPBC1 in contrast, we did not find iCAF but still there was a high population of myCAF that were intermixed with MP cancer cells (Suppl. Fig 4e, f). This result emphasizes the relevance of spatial technology to spatially contextualize the finding from snRNA-seq.

Representation of M2-polarized macrophages was observed across all bladder cancer samples analyzed by snRNA-seq (Fig. 3h). MPBC1 and MPBC2, in particular, exhibited a statistically significant higher M2-to-M1 macrophage (Fig 4g), as reflected by the expression of M2-associated markers (Fig 4g). Like CAFs, M2 macrophages were spatially integrated within the MP regions (Fig 4a), contributing to the intricate TME characteristic of this histology.

To investigate the TME in URO subtypes, we focused on spatially analyzing the oxygen-related features enriched in the BL subtypes. snRNA-seq results showed an enriched expression of “Hypoxia” and “Angiogenesis” pathways in the BL subtype as compared to the LLI and MP (Fig 3g). In addition, the immunohistochemical analysis showed that KRT5 positive basal cells were significantly closer to the vascular stroma (Fig 3k). Therefore, we analyzed the spatial distribution of the standard Hypoxia marker *EGLN3* within the BL component of MPBC1. *EGLN3* is a marker of sustained or late-stage hypoxic responses, and more stable than *HIF1A* mRNA, which can be transient (Cavadas et al. 2015). We observed strong overexpression of *EGLN3* in the BL component as compared with the MP histology (Fig 4 h,i) that was also validated in the snRNA-seq analysis of the same sample (Fig 4 j,k). This result was extended to the snRNA-seq 11 sample cohort showing that EGLN3 was higher in BL as compared to LLI and MP (Suppl. Fig 4g). We also analyzed the co-occurrence score of the endothelial populations relative to the BL and MP cancer cells. The results validated the spatial observation, showing that BL were more likely to co-occur with endothelial cells. (Fig 4l). These findings support that BL cells might exhibit a distinct sensitivity to oxygen availability, consistent with observations from immunohistochemical analyses. However, comparison with the LLI component was not possible due to its absence.

In summary, spatial transcriptomics complemented and spatially contextualized the snRNA-seq analysis, highlighting the complexity of the TME and revealing distinct differences between MP and URO histologies within the same tumor specimens.

## DISCUSSION

In this study, we define chromatin subtypes in HGT1 NMIBC based on enhancer profiling that show intratumor heterogeneity, are associated with activation of distinct molecular pathways, and with divergent interactions with the TME. Previous classifications using bulk RNA-seq lacked the resolution to uncover this complexity, (Sia Viborg Lindskrog et al. 2021) underestimating the significance of intratumor heterogeneity and limiting the ability to correlate molecular subtypes with patient’s clinical outcomes. More recently single-cell technologies have since unveiled the coexistence of subtypes within individual tumors (Sia V. Lindskrog et al. 2023b). However, further refinement and functional interpretation of these subtypes remained necessary. Our work builds upon these findings by anchoring subtype distinctions to their chromatin landscape. Specifically, our results provide chromatin support to UROMOL2021 subtypes 1, 2a, and 3. In contrast, LC subtype 2b (Sia Viborg Lindskrog et al. 2021), may represent a heterogeneous mixture of cellular states rather than a discrete, transcriptionally and epigenetically defined entity. This is supported by the lack of alignment between subtype 2b and any chromatin-defined subtype, as well as the absence of 2b signature scoring at the single-nucleus level in our cohort.

Our epigenetic profiling subdivides the URO tumors into two distinct chromatin subtypes, coming from potentially distinct cells of origin. While we cannot rule out the existence of additional subgroups, especially given the size limitation of our H3K27ac dataset, we show the presence of two major states: a luminal-like (LLI) and a basal-like (BL) chromatin program, sharing features with UROMOL2021 subtypes 2a and 4 respectively, respectively. These two states were found to coexist in approximately 40% of the tumors analyzed. Copy number variation (CNV) inference from snRNA-seq data revealed that both URO subtypes share similar genetic backgrounds within the same tumor. The immunohistochemical analysis further supported this coexistence of the classification showing that KRT5-positive basal-like cancer cells were frequently located near vascularized stroma, while KRT20-positive luminal-like cells tended to cluster toward the tumor center. Taken together, the non-genetic mechanism, spatial compartmentalization, and variable degrees of heterogeneity across cases suggest that the epigenetic subtypes may reflect tumor plasticity driven by environmental factors. For example oxygen accessibility has been previously implicated in cancer plasticity (Chakraborty et al. 2019) particularly through the oxygen-sensing activity of the H3K27 histone demethylase KDM6A. Interestingly, inactivating mutations in KDM6A, which are more common in bladder tumors with luminal features, may further support the role of hypoxia-mediated epigenetic reprogramming in shaping tumor heterogeneity.

In contrast, the two distinct histological subtypes, URO and MP, which can also coexist within the same tumor specimen, represent separate genetic clones and may correspond to more stable transcriptional states. Importantly, URO and MP exhibit markedly different interactions between cancer cells and their respective TMEs (TME). The MP subtype is characterized by a more complex microenvironment, with a diverse population of cancer-associated fibroblasts (CAFs) occupying spatially distinct regions. In MP, myofibroblastic CAFs (myCAFs) are found in close proximity to tumor cells, whereas inflammatory CAFs (iCAFs) are more intermingled with immune cells. This spatial organization coincides with a higher infiltration of M2-polarized tumor-associated macrophages (TAMs), within the MP component. Together, the presence of M2 macrophages and these CAF subtypes suggests an immunosuppressive microenvironment associated with the MP cellular niche. CAFs are increasingly recognized as key regulators of tumor progression through mechanisms such as extracellular matrix remodeling, secretion of growth factors, and modulation of immune responses (Sahai et al. 2020; Gascard and Tlsty 2016; Lavie et al. 2022). Similarly, M2-polarized TAMs promote tumor progression and immune evasion by supporting tissue repair, angiogenesis, and immunosuppression (Mantovani et al. 2017). CAFs can recruit and polarize macrophages toward the M2 phenotype(Chen et al. 2021) via cytokine and fibroblast growth factors (LeBleu and Kalluri 2018). The reciprocal interaction between CAFs and M2 macrophages reinforces an immunosuppressive TME and contributes to resistance to immune checkpoint inhibitors in multiple cancer types (Binnewies et al. 2018). The specific TME characteristics can contribute to the worse outcome associated with the MP histology. In contrast, the URO subtype exhibits an immune-excluded phenotype with fewer infiltrating immune cells, reflecting a less complex stromal landscape.

In summary, the presence of chromatin-based subtypes exhibiting intra-tumor heterogeneity and potential phenotypic plasticity underscores the complex challenges faced in cancer treatment. These potentially dynamic epigenetic states may enable tumors to adapt to therapeutic pressures, contributing to resistance and disease progression. Furthermore, the differences observed between micropapillary (MP) and urothelial (URO) subtypes are based on variations in the TME, suggesting that microenvironmental factors may actively drive or sustain divergent tumor programs. Together, these insights highlight the complex heterogeneity of bladder cancer and underscore the relevance of single-cell and spatial resolution analyses in uncovering disease biology, ultimately guiding the development of more precise and context-specific therapeutic strategies.

## Acknowledgement

P.C. acknowledges funding from the Ministry of Economy and Competitiveness, Instituto de Salud Carlos III (Institute of Health Carlos III)— PI23/01533. P.C is scientific advisor and co-founder of Cure51 biotech. JB acknowledges support from the Kaifer Grant.

## Author contributions

J.B., P.C., and H.W.L., conceptualized the study. S.K., M.G.M., S.L.V., A.F.T, M.L, performed experiments and reviewed the draft. S.H.L., M.L., N.J., S.M., contributed samples and performed histological analysis. I.E. analyzed clinical data. Y.X., X.Q., Y.J., M.N.D, R.L., performed computational analysis. P.C., J.B., H.W.L., M.B., T.C., provided supervision and oversight., P.C., J.B, H.W.L. wrote the manuscript.

## METHODS

### NMIBC tissues and FiTAc H3K27Ac profiling

The NMIBC tissues were obtained from collections at Hospital del Mar-Parc de Salut Mar-Biobank, Barcelona, Spain. For H3K27ac profiling we selected 17 NMIBC, 6 of them with identified micropapillary content. All the patients were treatment naive. To increase the enrichment in cancer cells, we macro-dissected FFPE sections whenever needed to obtain >80% tumor cells. To increase the enrichment in micropapillary content, cores were taken at areas of high MP content as selected by a pathologist.

The FiTAc-seq method was applied as previously described (Font-Tello et al. 2020). We started from 10 sections, 10 μm thick that were washed 3 times with xylenes to remove paraffin, rehydrated in an ethanol/water series. The tissue was resuspended in lysis buffer as previously described and sonicated for 5 minutes using a Covaris E220 instrument (setting: 140 peak incident power, 5% duty factor, and 200 cycles per burst) in 1 ml adaptive focused acoustics (AFA) fiber millitubes. Soluble chromatin (5 μg) was immunoprecipitated with 10 μg H3K27ac (Diagenode catalog number C15410196) antibody (Ab). ChIP-seq libraries were constructed using ThruPLEX-FD kits (Rubicon Genomics) following the manufacturer’s protocols. 75-bp single-end reads were sequenced on a Nextseq instrument (Illumina).

### Nuclei preparation and single cell ATACseq

For each of the two HGT1 cases, a 40 μm section was obtained and nuclei isolation was performed as previously described (Font-Tello et al. 2020; Cejas and Long 2022). In brief, the section was suspended in 300 μl of cold buffer comprising 0.1% NP40, 0.1% Tween-20, and 0.01% Digitonin. The homogenate tissue was then transferred to a pre-chilled 1.5 ml microfuge tube and incubated on ice for 10 min. Lysates were then filtered through a 40 μm cell strainer, and nuclei were centrifuged for 10 min at 1500 relative centrifugal force (RCF) in a pre-chilled (4°C) fixed-angle centrifuge. Nuclei were resuspended in 300 μl of buffer containing 0.1% Tween-20 and enumerated using a hemocytometer with Trypan blue stain. Approximately 7000 cells were targeted per sample and processed following the 10× Genomics scATAC-seq sample preparation protocol (Chromium Single Cell ATAC Library & Gel Bead Kit, 10× Genomics). 150 bp paired end reads were sequenced in a NovaSeq XP.

### Nuclei isolation and snRNA-seq from FFPE tissue

Nine NMIBC clinical samples were selected from the Hospital del Mar-Parc de Salut Mar-Biobank, Barcelona, Spain. Nuclei isolation was performed using a modified version of a protocol originally developed for frozen specimens (Cejas and Long, 2022), optimized for FFPE material.

Paraffin was removed through sequential xylene washes, followed by tissue rehydration in a graded ethanol series ending in distilled water. After centrifugation, the tissue pellet was resuspended in a dissociation buffer composed of phosphate-buffered saline (PBS, 896.5 µL), Liberase TM (38.5 µL at 6.5 U/µL), Collagenase D (40 µL at 100 mg/mL), and RNase inhibitor (25 µL at 40 U/µL). Samples were incubated at 37°C for 1 hour in a thermomixer to facilitate enzymatic tissue dissociation. Following dissociation, samples were washed with PBS, centrifuged, and the resulting pellet was resuspended in 300 µL of a lysis buffer containing 0.1% NP-40, 0.1% Tween-20, and 0.01% digitonin. The suspension was incubated overnight at 37°C in a thermomixer to ensure complete lysis and nuclear release. After incubation, 700 µL of 0.1% Tween-20 RSB buffer was added before passing them through a 40 µm cell strainer to remove debris. The nuclei could then be counted on the Countess II before proceeding to pelleting them down by centrifugation at 1,250 × g for 10 minutes at 4°C using a fixed-angle rotor.

Approximately 10,000 nuclei per sample were processed for single-nucleus RNA sequencing using the 10x Genomics GEM-X Flex Gene Expression protocol. Libraries were prepared, multiplexed, and sequenced as 150 bp paired end reads on the Illumina NovaSeq XP platform.

### Visium HD protocol

Spatial transcriptomic profiling was performed using the 10x Genomics Visium HD platform, following the manufacturer’s protocol with minor modifications. Formalin-fixed, paraffin-embedded (FFPE) tissue blocks were sectioned at 5 μm thickness and mounted onto standard glass slides. Sections were then fixed, stained with hematoxylin and eosin (H&E), and imaged using a high-resolution slide scanner to capture detailed tissue morphology. Following imaging, tissue sections underwent permeabilization to facilitate probe access. Subsequently, a probe hybridization step was carried out overnight to target mRNA sequences.

After hybridization, the slides were processed using the Visium CytAssist instrument. This step involved aligning the tissue sections on the standard glass slides with the capture areas on the Visium HD slide, enabling the transfer of spatially barcoded probes to the capture areas. The CytAssist instrument facilitates the transfer of spatial information from the tissue sections to the Visium HD slide. Library preparation was then completed using the Visium HD library construction kit, and sequencing was performed on an Illumina NovaSeq system to a depth sufficient for high-resolution spatial gene expression analysis.

### Immunohistochemistry

We performed immunohistochemistry on tissue microarrays made from a collection of 162 bladder tumor samples, composed mainly of HGT1 (102 cases) but also containing 41 low-grade tumors and 19 MIBC (Lloreta et al. 2017)). Tissue sections were deparaffinized in xylenes and hydrated through ethanol and water series. After antigen retrieval, slides were treated with 3% H2O2 in PBS for 10 min to quench endogenous peroxidases, washed, and incubated in blocking solution (PBS containing 1% BSA and 1% Tween-20) for 1h at ambient temperature. Slides were incubated with TP63(FLEX Monoclonal Mouse Anti-Human p63 Protein, VENTANA anti-p63 (4A4), Part Number:GA66261-2) KRT5 (KRT 5 VENTANA anti-Cytokeratin 5/6 (D5/16B4) Mouse Monoclonal Primary Antibody), KRT20 (KRT 20 VENTANA anti-Cytokeratin 20 (SP33) Rabbit Monoclonal Primary Antibody) (Ab diluted in blocking solution for 1 h. Slides were washed in PBS and incubated with the peroxidase-based EnVision Kit (Dako).

### Staining Scoring

Scoring has been performed by an expert pathologist. p63 stainings were assessed by Histoscore (H-score), a metric that considers both the percentage of stained cells and the staining intensity. In detail, H-score is calculated as follows: 3 x percentage of strongly staining nuclei + 2 x percentage of moderately staining nuclei + percentage of weakly staining nuclei. Intensity was scored: 0: no staining, 1: weak, 2: moderate, and 3: strong staining. For KRT 5/6 and KRT20, the percentage of positive cells was assessed.

### Computational and statistical analysis

#### H3K27Ac FiTAc-seq analysis

##### Alignment and peak calling

All samples were processed through the computational pipeline developed at the Dana-Farber Cancer Institute Center for Functional Cancer Epigenetics using primarily open-source programs (https://github.com/liulab-dfci/CHIPS)(Qiu et al. 2021). Sequence reads were aligned with Burrows-Wheeler Aligner (BWA) (H. Li and Durbin 2009) to build hg19 and uniquely mapped, non-redundant reads were retained. These reads were used to generate binding sites with Model-Based Analysis of ChIP-Seq 2 (MACS v2.1.1.20160309), with a q-value (FDR) threshold of 0.01(Zhang et al. 2008). We evaluated multiple quality control criteria based on alignment information and peak quality: (i) sequence quality score; (ii) uniquely mappable reads (reads that can only map to one location in the genome); (iii) uniquely mappable locations (locations that can only be mapped by at least one read); (iv) peak overlap with Velcro regions, a comprehensive set of locations – also called consensus signal artifact regions – in the genome that have anomalous, unstructured high signal or read counts in next-generation sequencing experiments independent of cell line and of type of experiment; (v) number of total peaks (the minimum required was 8,000); (vi) high-confidence peaks (the number of peaks that are tenfold enriched over background); (vii) Overlap with known DHS sites derived from the ENCODE Project (the minimum required was an 80% overlap); and (viii) peak conservation (a measure of sequence similarity across species based on the hypothesis that conserved sequences are more likely to be functional). Genome tracks were visualized by IGV (v2.14.1) (Robinson et al. 2017).

##### Unsupervised analysis of the H3K27ac dataset

We utilized Permutational Multivariate Analysis of Variance (PERMANOVA) to statistically assess differences among the three clusters identified through hierarchical clustering of gene expression data. PERMANOVA is a non-parametric method that evaluates whether the centroids of predefined groups differ significantly in a multivariate space. This analysis was conducted using the ‘Adonis2’ function from the ‘vegan’ package in R, which operates on distance matrices and employs permutation tests to determine statistical significance.

By applying PERMANOVA, we tested the null hypothesis that there are no differences in the multivariate centroids among the three clusters. The analysis was performed with 999 permutations to assess the significance of the observed differences. This approach provided a robust statistical framework to validate the clustering results and ensure that the observed groupings reflect meaningful biological variation rather than random chance.

##### Differential binding analyses

We used our COBRA pipeline for differential analysis (Qiu et al. 2021). Briefly, peaks from all samples were merged to create a union set of sites for each transcription factor and histone mark using bedops (Neph et al. 2012). Read densities were calculated for each peak for each sample and used for the comparison of cistromes across samples. Sample similarity was determined by hierarchical clustering using the Spearman correlation between samples with significant differences. PCA plots were generated using standard R tools. Differential peaks were identified by DEseq2 with adjusted P ≤ 0.05 and |log2FoldChange| > 0.5. A total number of reads in each sample was applied to the size factor in DEseq2, which can normalize the sequencing depth between samples. Peaks from each group were used for motif analysis by the motif search findMotifsGenome.pl in HOMER2 (v3.0.0)(Huppert et al. 2009), with cutoff q-value ≤ 1e-10. The signals of each sample on differential binding sites were visualized by Deeptools (Huppert et al. 2009; Ramírez et al. 2014). For the PCA of H3K27Ac signals in Fig1b, all peaks from H3K27Ac ChIP-seq data were used.

##### Cistrome Toolkit and GREAT analysis

We utilized the ‘cistrome toolkit’ web tool (R. Zheng et al. 2019; Mei et al. 2017) to predict which factors exhibit significant binding overlap with differential peaks between URO1 (LLI) and URO2 (BL). Differential peaks were also analyzed using the GREAT web tool (McLean et al. 2010) to predict functions. A threshold of single nearest genes within 400 kb was set. The prediction results for GO Biological Process were visualized using ggplot2 (Wickham 2009).

#### Bulk RNA-seq analysis

##### Quality control and differential analysis

Read alignment, quality control, and data analysis were performed using the Visualization Pipeline for RNA-seq (VIPER) (Cornwell et al. 2018). Alignment to the hg19 human genome was done using STAR v2.7.0f (Dobin et al. 2013) followed by transcript assembly using cufflinks v2.2.1 (Dobin et al. 2013; Trapnell et al. 2010) and RseQC v2.6.2 (Wang, Wang, and Li 2012). Differential gene expression analyses were performed comparing LLI to BL using DESeq2 v1.18.1 (Wang, Wang, and Li 2012; Mainz Mainz Press 2016), utilizing absolute gene counts for RNA-Seq data and raw read counts for transcriptomic profiling data. The samples in the LLI and BL groups were matched with samples from H3K27ac ChIP-seq analysis. Specifically, in the H3K27ac dataset, there were 6 samples assigned to URO1 (LLI) and 6 to URO2 (BL), while in the RNA-seq cohort, there were 3 samples for URO1 (LLI) and 4 for URO2 (BL). Gene Set Enrichment Analysis (GSEA) was conducted using the GSEA software (GSEA Java; v4.1.0) with Hallmark gene sets. Genes were pre-ranked based on Log2FC for the BL versus LLI comparison, and enrichment scores were computed (p = 1, weighted). The top 10 enriched gene sets on either side were visualized using ggplot2 (v3.5.1) (Subramanian et al. 2005).

##### Integration bulkRNA-seq with H3K27Ac ChIP-seq data

We visualized the differentially expressed genes from the RNA-seq analysis using a volcano plot. For the signatures of the LLI vs BL subtypes, we selected the top 15 differential genes from the RNA-seq analysis. These genes were chosen based on their correlation with the top differential peaks (ranked by adjusted P-value) identified in the corresponding H3K27ac ChIP-seq analysis.

##### Gene set variation analysis

The GSVA (Hänzelmann, Castelo, and Guinney 2013) score was computed for the entire RNA-seq cohort, and the disparities between the GSVA scores of LLI and BL were utilized for sample classification. Samples falling within the 1st quartile, reflecting the highest LLI - BL score, were classified as LLI-like, while those within the 4th quartile were categorized as BL-like. Boxplot representing markers’ expression in BL-like and LLI-like groups were plotted using ggplot2 v3.5.1. The GSVA analysis was also conducted on data from the UROMOL cohort. GSVA analysis was also performed on data from the UROMOL cohort. As described for the other cohort, samples in the highest quartile of the LLI–BL score distribution were classified as LLI-like, while those in the lowest quartile were designated as BL-like. For the UROMOL cohort, chi-square tests were conducted to assess associations between the four classes defined in the UROMOL2021 classification and the two subtypes identified by our LLI– BL classification.

##### bulkRNA-seq statistical analysis

Cox Proportional-Hazards analyses were done with the survival (v3.7.0) package and Kaplan-Meyer curves were plotted with ggsurvfit (v1.1.0). Markers’ expressions between LLI and BL cases were compared using a Student’s t test with Welch correction for the UROMOL cohort (n > 50, unequal variances - F test with p-value < 0.05) and a Wilcoxon rank-sum test for the CFCE cohort (n < 50, non-Gaussian distribution - Shapiro test with p-value < 0.05). All the statistical analyses were done using the ‘stats’ package (v4.3.2) in RStudio (R v4.3.2).

#### Single-cell ATAC-seq data processing

##### Quality control

The scATAC-seq data from two samples (URO1 - 7166 cells; URO2 - 1733 cells) underwent processing using the cellranger-atac (v2.0.0) (Satpathy et al. 2019) pipeline with default parameters. Quality control filtering of low-quality cells was performed using the R packages Seurat (v3) (Butler et al. 2018) and Signac (v1.6.0) (Butler et al. 2018; Stuart et al. 2021), based on criteria including nucleosome-binding pattern strength, transcription start site enrichment score, number of fragments in peaks>100,peaks > 600.

##### Sample integration

Integration of samples was achieved using a common peak set derived from peaks of each sample with peak widths ranging from 20 to 10,000. The peak-count matrix underwent TF-IDF normalization using the Signac package. Singular value decomposition (SVD), also referred to as latent semantic indexing (LSI), was performed on the normalized matrix using the RunSVD function in Signac, resulting in 2:30 LSI components. These components were then utilized for nonlinear dimensionality reduction via the RunUMAP function from the Seurat package. The Seurat function FindNeighbors was utilized to create a shared nearest neighbor graph based on the 2:30 LSI components. Subsequently, the FindClusters function was employed to iteratively cluster nuclei, optimizing modularity through the Louvain algorithm. The gene activity score for the integrated object was computed using the GeneActivity function and subsequently normalized using the NormalizeData function. Differential analysis for each cluster was conducted using the FindMarkers function, while differential motifs for each cluster were identified using chromVar. The LLI and BL signatures scores were computed using the AddModuleScore function. Motif enrichment analysis for each cluster was done by chromVAR (Schep et al. 2017).

##### Cell type annotation

The scATAnno (Y. Jiang et al. 2024) package was used to perform cell type annotation.

##### Single cell ATAC CNV inference

our internal CNV calling tool was used to call copy number variation from single-cell ATAC-seq samples, this tool adapted an existing bulk ATAC-seq method to utilize off-target scATAC-seq reads for inferring DNA copy number changes. The genome was divided into large intervals, and the coverage of each interval was determined. scATAC-CNV averaged the coverage of 100 GC-matched intervals to establish background levels. Comparisons between interval coverage and corresponding GC-matched background enabled estimation of CNV fold change. Interval sizes of 1–2 Mb were chosen to accommodate the sparse scATAC-seq data using the “ChunkGRanges” function in GenomicRange. The “GCcontent” function of biovizBase was used to calculate GC content for each interval. Coverage adjustment for removed peaks was achieved by incorporating the effective window size during calculation.

#### Fixed snRNA-seq (FLEX) data processing

##### Quality control, dimensionality reduction and clustering

Cellranger (version 6.0.2) (G. X. Y. Zheng et al. 2017) by 10x Genomics aligned reads to the prebuilt GRCh38 genome reference version 2020-A (refdata-gex-GRCh38-2020-A). R package Seurat was used to perform subsequent processing using the cell by gene matrix from cellranger. Doublets were identified and removed using the scDblFinder (Germain et al. 2021) package. Barcodes were filtered by UMIs, expressed genes, and mitochondrial gene percentage. Cell cycle genes and mitochondrial genes were regressed out using the function ScaleData. The filtered gene-count matrix underwent scaling and normalization for sequencing depth using Seurat’s ‘SCTransform’ function. Following this, principal components were calculated using Seurat’s RunPCA function. Cells were then clustered using Louvain graph-based approach. Initially, the Seurat function FindNeighbors was employed to construct a k-nearest neighbor graph based on Euclidean distances in principal component analysis (PCA) space. Cells with similar expression patterns were connected by edges. The first 30 principal components were used for this step with default parameters. Clustering was then performed using modularity optimization techniques, specifically the Louvain algorithm from the Seurat function FindClusters.

##### Data integration and cellt ype annotation

After quality control, nine filtered samples were merged. Subsequently, the merged objects were normalized using the Seurat SCTransform function, employing the same parameters as when normalizing individual objects. Cells were then clustered using the top 30 PCA dimensions via the FindNeighbors and FindClusters functions, with a Resolution parameter set to 0.2. Next, the RunUMAP function was applied to obtain new cell embeddings. The LLI and BL signature scores were calculated utilizing the AddModuleScore function. InferCNV (https://github.com/broadinstitute/inferCNV)(v1.19.1) was used to identify CNA in single cells with threshold: cutoff = 0.1, HMM = TRUE, leiden_method=“simple”, cluster_by_groups=TRUE, denoise=TRUE. For cell type annotation, we integrated the results from three sources: (i) outcomes derived from singleR(Aran et al. 2019) analysis, (ii) expression data of well-established marker genes, and(iii) results obtained from inferCNV analysis

##### Classification, GSEA

Tumor cells were classified by their signature scores. Cells were ranked by LLI and BL scores, and a quantile threshold was established. Cells in the 1st quantile of LLI but not meeting the BL threshold were classified as LLI-like. Similarly, cells meeting the BL but not the LLI threshold were classified as BL-like. Remaining cells were labeled as unidentified. The ssGSEA analysis was performed using ‘escape’ in scRepertoire toolkit for 3 classes with hallmark gene sets (Borcherding, Bormann, and Kraus 2020).

#### IHC image analysis

##### Alignment

Staining for KRT5 and KRT20 were done on 2 adjacent slides of the same FFPE block for each tumor. The plug-in StackReg (https://bigwww.epfl.ch/thevenaz/stackreg/) was used in FiJi (v1.53t) to align images. The same filters as previously described were used to detect KRT5 and KRT20 positive regions. Image annotations were done in RStudio (R v4.3.2).

##### Distances

19 images of tumors positive for both KRT5 and KRT20 (36% of the mixed tumors) were analyzed to measure the distance between KRT20-positive or KRT5-positive) cells and vascular stroma. Briefly, for each image, vascular stroma was outlined manually (by/under the supervision of) a patho-histologist using FiJi (v1.53t). Each colored image was then converted to black and white and stained areas were detected using the “Set Auto Threshold’’ tool with the “Moments’’algorithm. Because nuclei were stained with hematoxylin, they appeared in dark blue and were thus detected as part of the KRT5 or KRT20 positive areas. To distinguish the diaminobenzidine signal (marking the KRT20 or KRT5 cells) from the hematoxylin signal (marking nuclei), an additional filter was applied to remove from the KRT5+/KRT20+ regions the pixels that were more blue than brown. This was done in RStudio by removing pixels with a higher blue intensity than the red one. Finally, for each pixel in the stained area, the closest blood vessel pixel was identified using Python’s rtree library and the euclidean distance between them was measured. Images annotations and paired dot plots were done in RStudio (R v4.3.2). Wilcoxon rank-sum test was performed to compare distances between KRT5/KRT20 positive pixels and vascular stroma.

#### Spatial transcriptomics data processing and analysis

Raw sequencing reads were processed using the 10x Genomics Space Ranger HD pipeline (version 3.1.3), which included alignment to the GRCh38 human reference genome and assignment of reads to spatial barcodes. Gene expression matrices were generated at 2-µm resolution. Cell segmentation was performed with bin2cell(Polański et al. 2024), using the binned output from Space Ranger as input, with parameters optimized for spatial resolution.

Segmented cell-level gene expression data were analyzed with Scanpy (version 1.10.2)(Wolf, Angerer, and Theis 2018). Low-quality cells and lowly expressed genes were filtered based on gene count, UMI count, and mitochondrial RNA content. Data were normalized, log-transformed and scaled. Highly variable genes were selected using the seurat_v3 method. Principal component analysis was performed, followed by neighborhood graph construction, UMAP dimensionality reduction, and Leiden clustering to identify transcriptionally distinct cell populations.

##### Cell type annotation for spatial transcriptomics

Cell types were assigned by integrating automated reference-based predictions from CellTypist(Domínguez Conde et al. 2022) with manual curation based on differentially expressed marker genes from Leiden clusters. Marker genes were identified via differential expression analysis, and cell identities were confirmed using known lineage markers and tissue context.

##### CAF subtype classification and immune signature scoring

Cancer-associated fibroblast subtypes were classified by expression of FN1 (myCAFs), C3 (iCAFs) and CD74 (apCAFs). Spatial co-occurrence between CAFs and other cell types was quantified using the gr.co_occurrence() function from the Squidpy package. Macrophage polarization was assessed by calculating M1 and M2 signature scores with the tl.score_genes() function in Scanpy, based on established gene sets.

##### Spatial transcriptomics Visualization

Data visualization was performed using Scanpy and Squidpy(Wolf, Angerer, and Theis 2018; Palla et al. 2022) plotting functions, including UMAP embeddings, spatial feature plots, and co-occurrence heatmaps to illustrate cellular distribution and spatial interactions.

