## Supplementary material for "Epigenetic Subtypes of High-Grade T1 Bladder Cancer Reveal Intra-Tumor Heterogeneity and Distinct Interactions with Tumor Microenvironment": Suppl. Information

Supplementary Figure 1

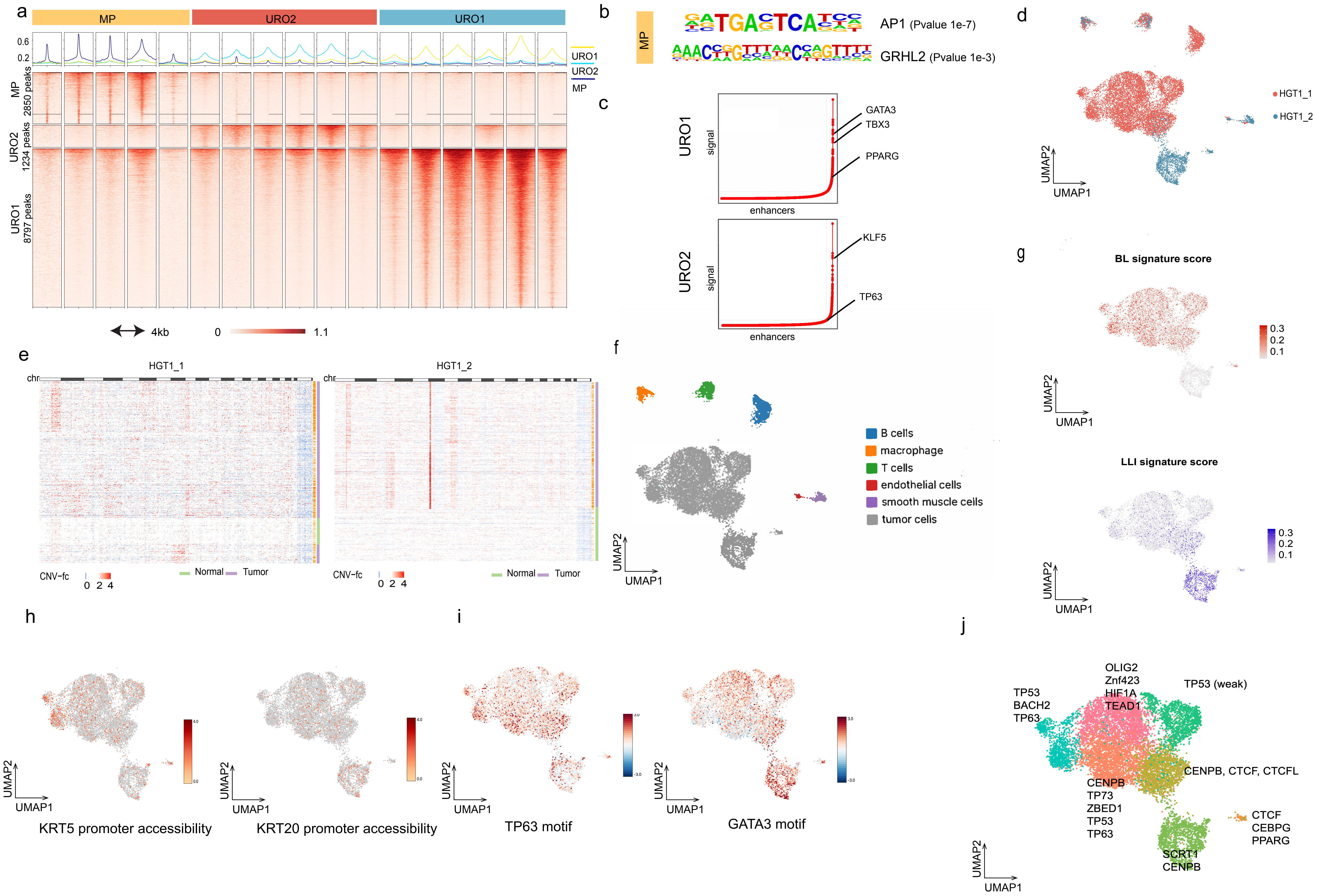

**Supplementary Figure 1**

**a)** Heatmap representation of the differential H3K27ac regions distinguishing MP, URO2 and URO1 clusters. **b)** Results from motif analysis at the MP-specific differential peaks. **c)** Signal distribution of H3K27ac marked enhancers from representative cases of a URO1 sample and a URO2 sample. Indicated are relevant super enhancers identified by the ROSE algorithm. **d)** UMAP analysis combining the two scATAC-seq datasets (HGT1_1 and HGT1_2) colored by case. **e)** CNV analysis of the tumor cells for the 2 scATAC-seq samples (HGT_1 and HGT_2). Rows are individual cells that have been clustered with K-means. Green bar indicates normal cells, purple bar indicates tumor cells. **f)** Annotation of the normal cell clusters in the scATAC-seq data (Methods). **g)** Combined UMAP analysis of HGT1_1 and HGT1_2 showing the gene activity score enrichment of the BL (red, top) and LLI (blue, bottom) signatures. **h)** Combined UMAP analysis of HGT1_1 and HGT1_2 showing accessibility at KRT5 and KRT20 promoters and **i)** enrichment of TP63 and GATA3 motifs. **j)** Combined UMAP analysis of HGT1_1 and HGT1_2 showing enriched motifs at the clusters defined by the Louvain graph algorithm (as described in Methods).

Supplementary Figure 2

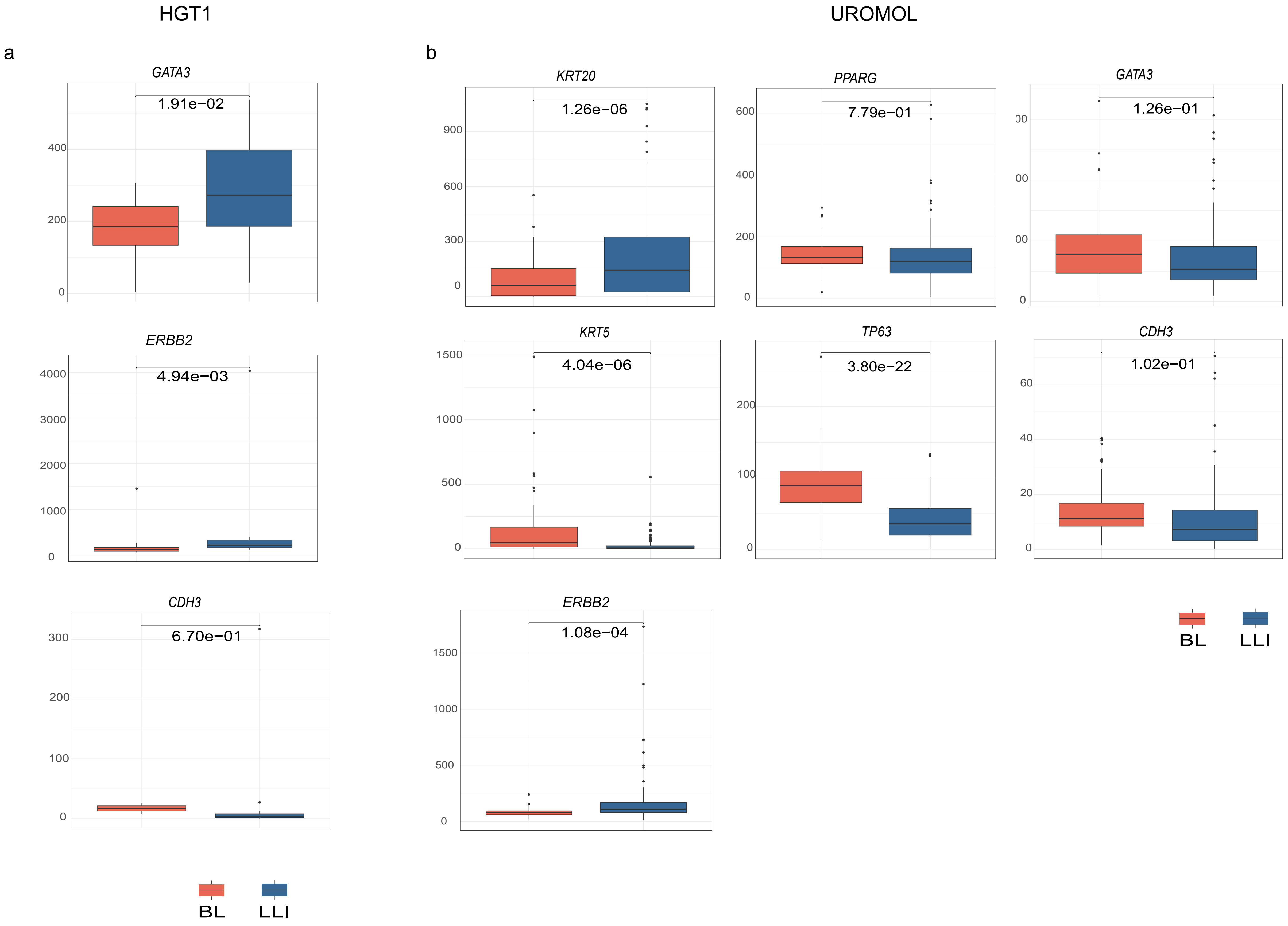

**Supplementary Figure 2**

**a)** Boxplots comparing the expression of marker genes analyzed in cases scored as BL (red) or LLI (blue) in the HGT1 cohort. The two groups are defined as the top high and top low quartiles and analyzed by t-test. **b)** Boxplots comparing the expression of marker genes analyzed in cases scored as BL (red) or LLI (blue) in UROMOL cohort. The two groups are defined as the top high and top low quartiles and analyzed by t-test.

Supplementary Figure 3

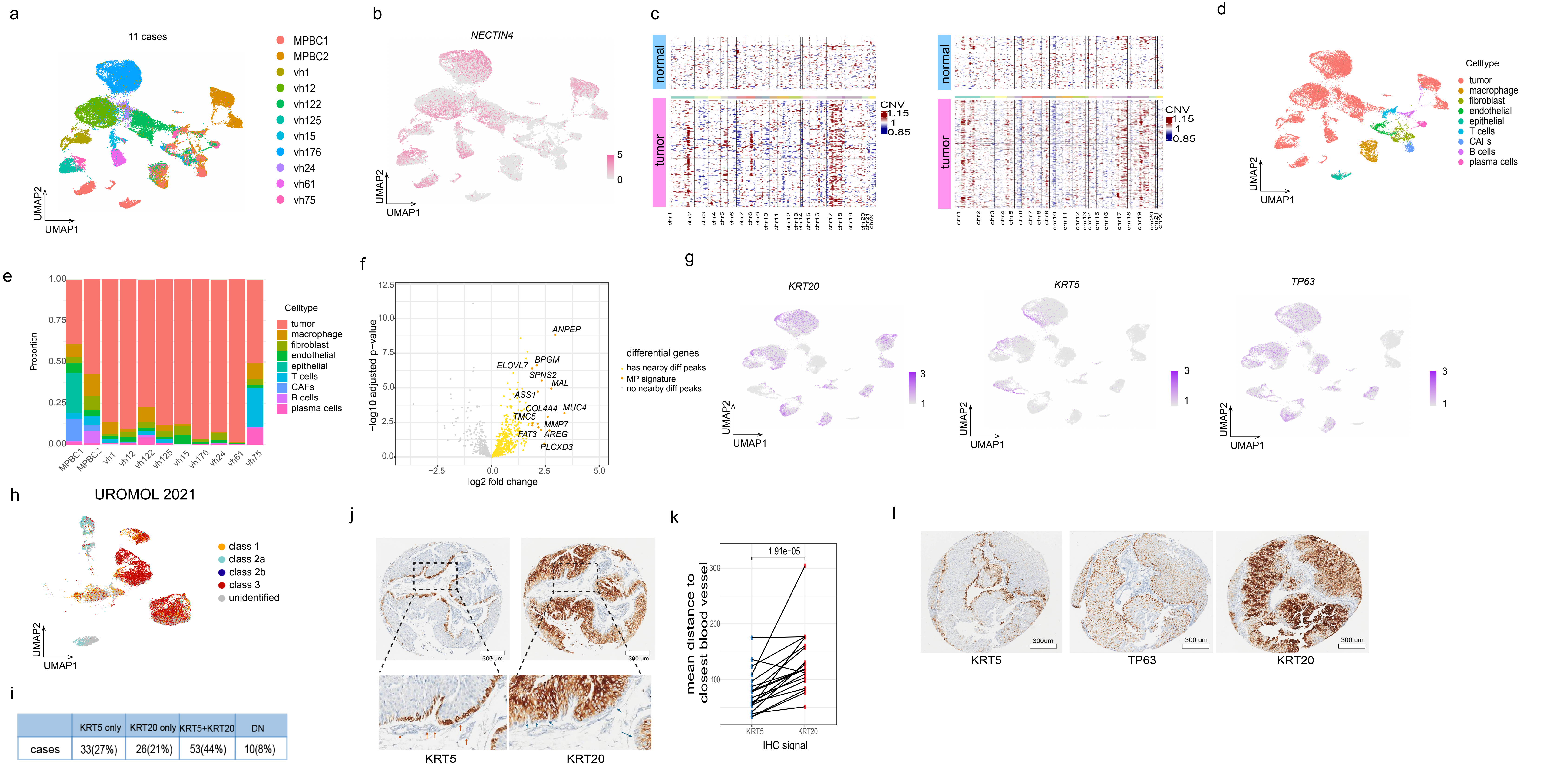


| | KRT5 only | KRT20 only | KRT5+KRT20 | DN |
|---|---|---|---|---|
| cases | 33(27%) | 26(21%) | 53(44%) | 10(8%) |

**Supplementary Figure 3**

**a)** UMAP analysis of all eleven snRNA-seq datasets that produced 37,879 nuclei colored by sample ID. **b)** UMAP analysis showing expression of *NECTIN4*. **c)** CNV analysis of two representative tumors analyzed by snRNA-seq (vh75 (top) and vh125 (Bottom)). **d)** UMAP analysis of all eleven snRNA-seq datasets annotated by cell type. **e)** Proportional barplot showing the relative percentage of each cell type in each tumor sample. **f)** Association between differential H3K27ac regions and differential gene expression for MP samples. The volcano plot depicts RNA-seq log2-fold change (x-axis) and p-value adjusted for multiple hypothesis testing (y-axis) as calculated by DESeq2. Each volcano plot depicts RNA-seq log2-fold change (x-axis) and p-value adjusted for multiple hypothesis testing (y-axis) as calculated by DESeq2. Each dot represents one gene, with yellow dots (left) indicating significant differentially expressed genes (DEG) associated with a differential H3K27ac region nearby for the MP. Gray dot: DEGs with no significant differential H3K27ac region nearby. **g)** UMAP analysis of all eleven snRNA-seq datasets showing gene expression for *KRT20* (upper), *KRT5* (middle) and *TP63* (lower). **h)** UMAP analysis of the cancer cells from the nine HGT1 tumors, showing scoring for the UROMOL2021 classification. Cells are assigned to one of the four UROMOL2021 classes: class 1 in orange, class 2a in light blue, class 2b in dark blue and class 3 in red, or to unclassified cells in grey. No cells classified as class 2b were detected at the single cell level. **i)** Summary of IHC scoring for KRT5 and KRT20 staining of the TMA, tumors are classified as KRT5 positive, KRT20 positive, KRT5 and KRT20 positives or KRT5 and KRT20 double negative (DN). **j)** Representative staining for KRT5 and KRT20 showing the distinct spatial locations of these stainings. The expanded view illustrates the anticorrelation of KRT5 and KRT20 staining, also indicating the proximity of the KRT5 positive cells to blood vessels (red arrows) and KRT20 negative (blue arrows) to blood vessels. **k)** Comparison of the mean distance to the nearest blood vessel for the KRT5 positive and KRT20 positive cells for cases showing positivity for both stains (N=16). **l)** Staining of a representative case for KRT5, KRT20, and TP63 shows that TP63-high cells exhibit greater overlap with KRT5 expression than with KRT20.

Supplementary Figure 4

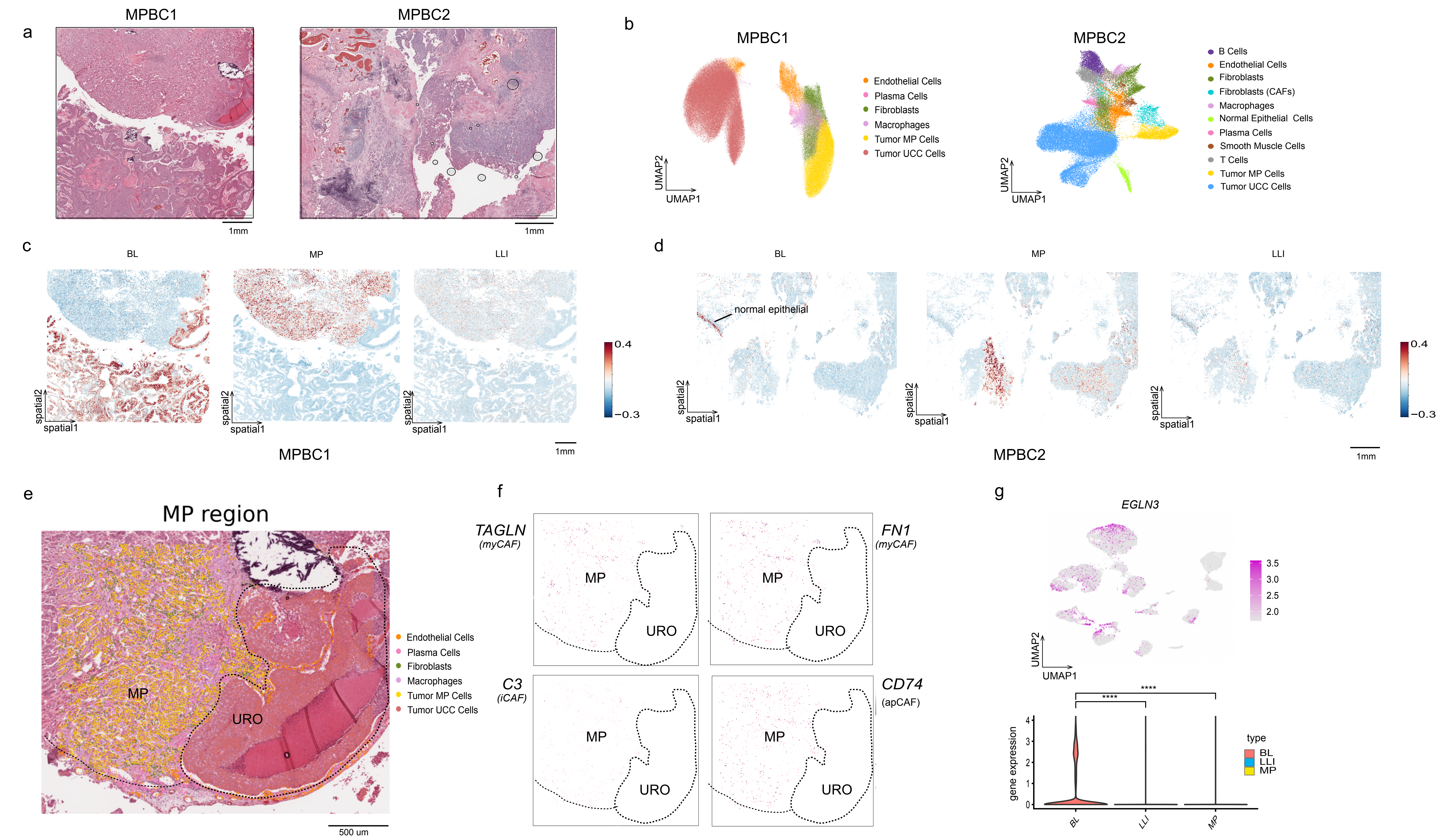

**Supplementary Figure 4**

**a)** H&E staining of the two MPBC cases analyzed by spatial transcriptomics. **b)** UMAP plot showing clustering by cell type for MPBC1 (left) and MPBC2 (right). **c)** Spatial representation of the scoring of all the cells by the CDS classification system as BL (left) MP (middle) and LLI (right) MPBC1. **d)** Spatial representation of the scoring of all the cells by the CDS classification system as BL (left) MP (middle) and LLI (right) MPBC2. **e)** Spatial mapping of CAFs in MPBC1 reveals distinct distributions between the MP and URO components. The MP region (dashed area, left) contains a high density of CAFs interspersed with tumor cells, whereas the URO region (dashed area, right) shows minimal CAF presence. **f)** spatial representation of canonical CAF markers in the MP and URO region. Results show the reduction of CAFs in the URO region. **g)** Top: UMAP analysis showing expression of *EGLN3* across cancer cells from the eleven snRNA-seq datasets. Bottom: violin plot displaying *EGLN3* expression in snRNA-seq cells scored as BL, LLI and MP. EGLN3 expression is significantly elevated in BL cells compared to the other groups ($p < 0.001$, t-test).
